# A Novel Chromatin-Opening Element for Stable Long-term Transgene Expression

**DOI:** 10.1101/626713

**Authors:** Shireen S. Rudina, Christina D. Smolke

## Abstract

Long-term stable expression of transgenes in mammalian cells is a challenge in gene therapy, recombinant protein production, and mammalian synthetic biology due to epigenetic silencing and position effect variegation. While multiple classes of regulatory elements have been discovered and proposed to help stabilize expression, the most efficacious has been the Ubiquitous Chromatin Opening Element (UCOE), and in particular, the prototypical A2UCOE from the *HNRPA2B1-CBX3* locus. We developed a feature-driven bioinformatics algorithm to discover other putative UCOEs from the human genome, and identified a novel UCOE (SRF-UCOE) that can resist transgene silencing in the methylation-prone P19 cell line. We demonstrate that a 767 bp core sequence of SRF-UCOE is modular to four common mammalian promoters. Notably, SRF-UCOE stabilizes gene expression in transduced P19 cells up to 2.4-fold better over 26 days than the existing A2UCOE by resisting constructs’ susceptibility to DNA methylation and histone deacetylation. Unlike existing UCOEs, SRF-UCOE lacks inherent transcriptional initiation activity, which can bolster its safe and predictable use in gene therapy constructs. We expect that expanding the set of UCOEs available will expand their utility to novel applications in gene therapy, synthetic biology, and biomanufacturing, as well as contribute to understanding their molecular mechanism.

## INTRODUCTION

Despite increasing sophistication in genome editing and transgene expression, stable long-term transgene expression in mammalian cells remains a bottleneck to the most important applications of heterologous gene expression. Biomanufacturing of therapeutic proteins requires the costly and inefficient isolation of rare, clonal cell lines that can stably express the transgene, as most clones show highly variable transgene expression and a susceptibility to time-dependent silencing of the gene (1,2). Similarly, most gene and cell therapies rely on the stable incorporation and expression of therapeutic genetic material, but are plagued by transgene expression variability and silencing, reducing the treatment efficacy (3). This loss of expression has been attributed to epigenetic silencing mechanisms, including DNA methylation, histone deacetylation, and chromatin condensation at the integration site (3,4). Random integration methods, such as retroviral vectors, amplify this effect due to the transgene’s interaction with its immediate genetic neighborhood, which may be heterochromatic (5).

Regulatory elements that address the problem of transgene variegation and silencing to confer long-term expression have traditionally fallen into two categories: passive boundary elements and active chromatin remodelling elements (6). The most widely used passive boundary element is the chicken **β**-globin 5’HS4 (cHS4) element, a traditional enhancer-blocking insulator that also functions as a barrier to heterochromatin spreading (7). In some applications, cHS4 is used to counteract position effects and can confer some stability to transgenes compared to the lack of an insulator (8,9). However, cHS4 and other passive insulators like Matrix Attachment Regions (MARs) can be cumbersome to use because one copy of the element is required on each side of the genetic construct, reducing available space for the transgene and any other desired genetic elements. Additionally, the element is highly cell type dependent, with limited utility in non-blood cell lineages (10,11). In contrast, active chromatin remodelling elements like ubiquitous chromatin opening elements (UCOEs) have gained popularity in the last decade because of their increased efficacy in resisting silencing (12).

UCOEs have been defined by their ability to confer reproducible, stable expression of transgenes, even when integrated into centromeric heterochromatin (13). One particular UCOE sequence from the *HNRPA2B1-CBX3* locus (dubbed A2UCOE) is by far the most studied and utilized of the currently identified UCOEs. The A2UCOE element encompasses a methylation-free CpG island between the *HNRPA2B1* and *CBX3* housekeeping genes. Stable expression can be achieved by using its innate promoter for *HNRPA2B1*, or as a regulatory element linked to any heterologous promoter (Figure 1a). Its efficacy has been attributed to its resistance to DNA methylation-mediated silencing and recruitment of chromatin remodellers (14). A2UCOE has demonstrated its utility in conferring long-term stable expression to gene therapy constructs in a variety of cell types and tissues, both *in vitro* and *in vivo* (15–17), and even in clinically-relevant human induced pluripotent stem cells (iPSCs) (18). Additionally, A2UCOE has demonstrated utility in the rapid selection and isolation of highly expressing clones in biomanufacturing to significantly improve titers (12,19,20). Recently, synthetic biologists have used the A2UCOE sequence to confer stability in dCas9-effector platform cell lines for CRISPR interference (CRISPRi) screens that perturb specific genes to study biological phenomena (21,22).

**Figure 1.**
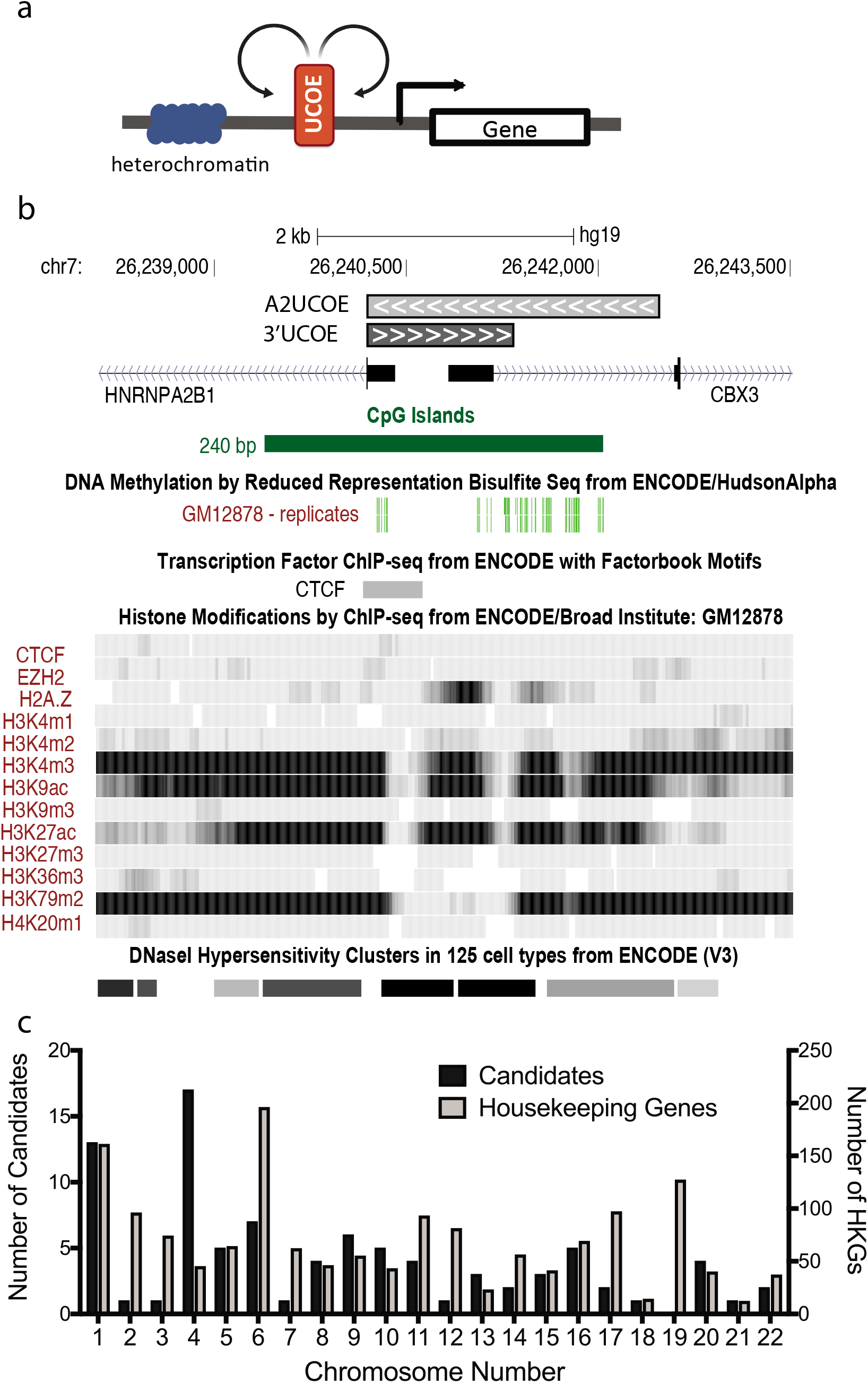
Computational screen based on characterized features of A2UCOE. a) Ubiquitous chromatin opening elements (UCOEs) are positioned directly upstream of a promoter driving expression of a transgene of interest. UCOEs function by actively maintaining an open chromatin structure such that transcription factors and RNA polymerases can interact with the promoter and drive transgene expression. b) Adapted UCSC Genome Browser screenshot of the *HNRPA2B1-CBX3* locus on chromosome 7 of the human genome (hg19 assembly). The locus consists of an unmethylated CpG island spanning the regulatory region between the two divergent housekeeping genes. Data utilized in screen, including Broad Institute/ENCODE ChIP-seq, reduced bisulfite sequencing (ENCODE), and transcription factor ChIP-seq for CTCF are shown. Also shown are DNAse hypersensitive sites as a measure of the openness of the locus. A2UCOE is a 2.2 kb region in the orientation of the *RPA2B1* gene, while 3’UCOE is a 1.2 kb region in the orientation of the *CBX3* gene. c) Distributions of the 1,522 housekeeping genes (HKG) identified in (30) and the 87 candidate UCOE results from the bioinformatics algorithm across the 22 human autosomes.

However, the efficacy of the A2UCOE element varies depending on promoter choice and cell type. Initial studies identified the entire 8 kb locus, from which it was shown that a 2.2 kb A2UCOE region maintained anti-silencing activity (13). Soon after, the 1.5 kb core region was identified, along with the discovery that a 1.2 kb sub-fragment of that element was equally effective in promoting stable expression (14,17). Since then, a variety of individual studies have found efficacy from variable lengths of the core sequence, but those additional truncations, up to 600 bp, have not been proven out further than the one specific application in each study (16,23,24). Thus, there remains a need for a single modular sequence under 1 kb that can predictably stabilize a broad diversity of gene expression constructs. Also, although A2UCOE seems to maintain the specificity of tissue-specific promoters, there is still a concern that its internal bidirectional promoter can cause non-specific transcriptional activation upon integration, and these off-target effects have traditionally been a concern in gene therapy (25). Additional UCOEs may be able to address the deficiencies of A2UCOE to find utility in more applications, as well as elucidate the underlying mechanism of this interesting class of elements.

With A2UCOE’s broad utility in mind, we sought to discover additional UCOE elements from the human genome that could complement or outperform A2UCOE. We designed a computational algorithm to search the human genome for other putative UCOEs and screened a dozen of the resulting sequences. Ultimately, we discovered and characterized a novel UCOE, named SRF-UCOE, which across four different promoters, stabilized expression in transduced P19 cells as well as or up to 2.4-fold better than the 2.2 kb A2UCOE and its 1.2 kb reverse complement (3’UCOE). Most notably, we found that SRF-UCOE does not have inherent transcriptional activity, which we posit will be an advantage for gene therapy applications where the bidirectional promoter nature of A2UCOE may cause unintended effects.

## MATERIAL AND METHODS

### Computational algorithm for identifying putative UCOE elements in the human genome

Data for the hg19 assembly of the human genome was downloaded from the appropriate sources, including (a) Broad Institute ChIP-seq data for GM12878 cells as part of the ENCODE consortium (13 tracks) (26), (b) UCSC Known Genes Track (27), (c) UCSC Genome Browser CpG Island track (further described in Supplementary Text), (d) ENCODE Reduced Representation Bisulfite Sequencing (RRBS) in GM12878 (28), and (e) the ENCODE transcription factor ChIP dataset (29).

Briefly, the 13 ChIP-seq tracks (consisting mostly of 11 histone modifications) for GM12878 cells were combined in a binary fashion (present or absent) to return a list of regions that contained the same combination of features. These sequences were then screened to remove regions that fell completely within a gene’s coding sequence according the UCSC Known Genes track. Next, sequences that did not consist of at least 50% overlap with a CpG island were removed, and the remaining sequences were screened for <20% methylated reads through RBSS data. Finally, regions were screened for a verified CTCF binding site. More specific details in Supplementary Text, and code is available at Github [www.github.com/srudina/UCOEdiscovery].

The ranking method was powered by data from a study identifying 1,522 housekeeping genes and their coefficient of variation across 42 tissues (30). Results from the computational algorithm were ranked first by their distance to the transcription start site of the nearest housekeeping gene, and then by the coefficient of variation of that gene.

### Construction of vectors

Actual candidate sequence was determined by a visual inspection of the outputs of the algorithm using the Feb 2009 GRCh37/hg19 assembly in the UCSC Genome Browser (http://genome.ucsc.edu/) (31). Start and end positions were visually determined for the candidates based on including as many desired features from the computational search as possible (e.g., the entirety of a CpG island, or to include any nearby CTCF binding sites) to result in a 1-1.5 kb length (Supplementary Table 2).

Primers were designed using NCBI Primer-Blast (Supplementary Table 5) and the candidates were PCR amplified from a U2OS (human bone osteosarcoma cell line) genome prep as template with the Kapa Hifi Hotstart Polymerase (Roche) according to manufacturer’s instructions. The primary stable transfection plasmid pCS4255 was created through the addition of the back-to-back EF1α-EGFP, hPGK-PuroR cassette using the Sal1/BglII sites in the ROSA26 donor plasmid, a gift from Charles Gersbach (Addgene plasmid #37200). Positive controls (i.e., 2.2 kb A2UCOE and 1.2 kb 3’UCOE elements) and putative UCOEs were cloned through ligation cloning into the Sal1/Nhe1 restriction enzyme sites in pCS4255. All plasmids from this study are shown in Supplementary Table 4.

Lentiviral vectors were based on the donor plasmid pCS3799. The UCOE-EF1α-EGFP cassette from the stable transfection plasmids was cloned into the Xma1/Xba1 sites in pCS3799 to make pCS4276. Additional truncation candidates were cloned into the Sal1/Nhe1 sites preceding EF1α. The three other promoters – cytomegalovirus (CMV), human phosphoglycerate kinase 1 (PGK), and respiratory syncytial virus (RSV) (sequences in Supplementary Table 6) - were cloned using the Sal1/Age1 sites in pCS4276, and UCOE candidates were cloned through the Sal1/Nhe1 sites in these plasmids.

### Maintenance of P19 cell lines

Mouse embryonic teratocarcinoma stem P19 cells were obtained from ATCC (CRL-1825) and maintained in alphaMem medium with Glutamax (Thermo Fisher Scientific) and 10% FBS (Thermo Fisher Scientific). Cells that were FACS-sorted were maintained in this growth media with the addition of 1% penicillin/streptomycin (Thermo Fisher Scientific). HEK293T (ATCC CRL-3216) cells for lentiviral production were cultured in DMEM media (Thermo Fisher Scientific) supplemented with 10% FBS (Thermo Fisher Scientific). All cells were grown at 37°C, 5% CO_2_, and 80% humidity in an incubator.

### Stable transfection of P19 cell line

P19 cells were seeded at 25,000 cells/well in 12-well plates. 24 h after seeding, cells were transfected using Lipofectamine 2000 (Thermo Fisher) according to the manufacturer’s instructions using 500 ng DNA/well and 2.5 μL of lipofectamine per well. 24 h after transfection, selection was initiated with 1 μg/mL puromycin (Sigma-Aldrich) in regular growth media, and from then on, cells were passaged at 1:15 or 1:20 dilutions whenever cells were 80-90% confluent (every 2-3 days), with frequent refreshing of puromycin-containing media to clear dead cells. After approximately 14-16 days, remaining cells were assumed to be stably transfected and were changed to regular growth media to initiate silencing experiment. During silencing experiments, the P19 cells were passaged every 2-3 days and re-seeded at 15,000 cells/well.

### Lentiviral preparation & transduction of P19 cell line

Two plasmids (pCS3800, encoding HIV-1 Gag; pCS380, encoding VSVg envelope protein) were used with varying versions of the donor plasmid pCS3799. HEK293T cells were plated at 5 x 10^6 cells in a 10 cm dish. 24 h after plating, the three plasmids (10 μg donor, 8 μg pCS3800, 10 μg pCS3801) were co-transfected together using a calcium phosphate protocol (32). The total DNA was brought to 500 μL in water, to which 500 μL of 2X HEPES-buffered saline, pH 7.0 (Alfa Aesar) was added and mixed. One-tenth of the total volume (100μL) of 2.5 M calcium chloride (Sigma-Aldrich) was then added to the mixture, followed by a 20-min incubation. The mixture was then added to the plate in a dropwise manner. Media was replaced 6 h later, and the supernatant was collected at 48 h after the transfection, filtered through 0.4 μm filter, and frozen at −80°C in 1 mL aliquots. Lentiviral aliquots were thawed in a 37°C bead bath before transductions.

P19 cells were plated at 20-30,000 cells/well in 12-well plates 1 day before transduction. 24 h after plating, cells were transduced at varying dilutions of the lentiviral stock (ranging from 1:2 to 1:100) in DMEM + 10% FBS with 8 μg/mL polybrene (Santa Cruz Biotechnologies). Media was refreshed on P19 cells 24 h after transduction, and cells were passaged and assayed through the Miltenyi VYB (see flow cytometry methods) 48 h after transduction. MOI was determined by reporter expression at this timepoint using the following formula:

MOI = ln(1/1-p) where p is the % of cells that are GFP positive at 48h post-transduction. (33)

Only populations that resulted in MOIs between 0.15 and 0.5 were subject to FACS sorting 5 days post transduction. Cells were FACS-sorted on the BD Influx cell sorter at the Stanford FACS Facility using the 488-nm laser and 525/40 filter to assay GFP expression. After gating for singlets and viability, GFP+ gate was drawn to be <0.1% GFP positive in non-transduced P19 cells. GFP+ gates were re-drawn for each promoter set to avoid the ~10% highest and lowest expressing cells within the GFP+ gate, but the same gate was used for every experimental condition under the same promoter. Triplicate wells of each population (12-15,000 cells/well in a 24-well plate) were collected.

### Epigenetic effector experiments

Replica-plated cell populations were treated with varying concentrations of 5-aza-2’-deoxycytidine (Sigma-Aldrich, A3656) or Trichostatin A (Sigma-Aldrich, T1952) 24 h after passaging. TSA was purchased as a readymade 5 mM solution in DMSO, which was then diluted to a 0.05 μM or 0.1 μM concentration in P19 growth media. 5 mg of 5-aza-2’-deoxycytidine was dissolved in 1 mL of 1:1 acetic acid:water to make a 21.9 mM stock solution, which was then diluted in P19 growth media to 2 μM or 10 μM. Cells were assayed through flow cytometry after 24 h.

### Flow cytometry analysis

Fluorescence data throughout silencing experiments was obtained using a MACSQuant VYB flow cytometer (Miltenyi Biotec). EGFP was measured through the 488-nm laser and 525/50nm band pass filter. Flow cytometry data was analyzed using the FlowJo software (Tree Star). After being gated for singlets and viability, GFP+ gates in Flowjo were drawn such that a non-transfected or non-transduced P19 cell population was at 1% GFP+. Median values reported are of cells within the GFP+ gate. Both % GFP positive and median are reported with the standard deviation of biological replicates.

## RESULTS

### Bioinformatics algorithm identifies 87 candidate UCOE regions

In order to develop criteria for identifying potential UCOE elements, particular properties of the A2UCOE locus in the human genome were identified that have been hypothesized to contribute to its mechanism (Figure 1b). A2UCOE encompasses divergently transcribed promoters of the *HNRPA2B1* and *CBX3* housekeeping genes, including a methylation-free CpG island. Distinct histone modification patterns, especially H3 and H4 acetylation, as well as the H3K4me3 mark that is associated with active transcription, have also been studied at this locus (34). Finally, insulator factor CTCF is known to bind to boundary regions and mediate three-dimensional chromatin loops at epigenetically distinct boundaries, making CTCF binding sites a hallmark of insulators (35). With the exact mechanism of A2UCOE’s functionality unknown, we chose to do as broad of a feature search as possible, hypothesizing that there are likely many other sequences in the human genome that perform similarly.

We examined the human genome through an algorithm that identifies areas with similar features to the A2UCOE locus. Because it is a chromatin-remodelling element, we naturally turned to the epigenetic signature at the locus as the first indicator of UCOE activity. With the causal effect of most histone marks still unknown (36), we chose to do as unbiased of a search as possible, by using all 13 of the ChIP-Seq tracks available to us through the Broad Institute/ENCODE consortium for what we determined to be the most karyotypically-normal somatic cell line, the GM12878 lymphoblastoid cell line. We searched for regions with the same pattern of presence/absence of histone marks (as well as three other DNA-associated proteins, EZH2, H2AZ, and CTCF) as measured by ChIP-seq across the hg19/Gr37 human genome assembly. This search resulted in 2,911 candidate regions. Since the sequence is a regulatory sequence, we further queried these regions to ensure that they did not fall completely within the coding sequence of genes, using the UCSC Known Genes track. Applying this filter reduced the candidate list to 936 regions. Next, based on 84% overlap of A2UCOE and the CPG island between *HRNPA2B1* and *CBX3*, we applied a condition that the region is largely composed of a CpG island. Specifically, we required regions to have at least a 50% overlap with a CpG island, bringing the number of candidate regions to 151. To ensure that we identified regions with unmethylated CpG islands, we further selected candidate regions based on Reduced Representation Bisulfite Sequencing (RRBS) data of GM12878 cells, also from the ENCODE project (further described in Supplementary text). The application of this criteria reduced the candidate list to 94 unique regions of the genome. As a final filter, we confirmed the CTCF binding sites with a different dataset from ENCODE, the ENCODE Transcription factor ChIP dataset, which encompassed data across several cell lines (29), bringing the number of candidate regions down to 88. The candidate list includes the A2UCOE locus on chromosome 7, and sizes of the candidate regions ranged between 57 to 3916 bp (Supplementary Figure 1).

To better prioritize the resulting candidate UCOEs for experimental characterization, we implemented a ranking methodology based on our hypothesis that the best UCOEs would be colocalized with the strongest housekeeping genes. We utilized a study of the human genome that identified 1,522 housekeeping genes using the gene expression profiles of 42 tissues (30). Elements were ranked first on the distance to the nearest housekeeping gene, and then by the coefficient of variance of that housekeeping gene according to that study. As a validation of our approach, the region encompassing the A2UCOE locus ranked first with this methodology, leaving 87 other ranked candidate regions to test for UCOE activity (Supplementary Table 1).

As many of the criteria used are broadly associated with regulatory regions of housekeeping genes, the distribution of the candidates across 22 autosomes was compared to the distribution of known housekeeping genes (Figure 1c). The results from this analysis show that the distributions are not correlated. For example, there are no candidates on chromosome 19 even though it has the third-most housekeeping genes, and chromosome 4 is overrepresented in the candidate UCOE regions compared to the distribution of housekeeping genes. The difference in distributions suggests that the properties the algorithm selects for are not uniformly distributed across housekeeping regulatory areas, and supports the utility of our algorithm.

The first ten candidate regions were visually inspected in the UCSC Genome Browser (31) in the hg19 assembly to draw candidate element boundaries such that the size of all tested candidates is between 1-1.5 kb (Supplementary Table 2). Boundaries were drawn to most conservatively include all nearby CTCF sites and the entirety of the CpG island when possible. Candidate regions were oriented in the same 5’ to 3’ direction as the nearest gene. In areas between dual divergent genes (i.e., Candidates 1, 6, 9, and 10), candidates were tested in both configurations with the (-) strand designated as “R”.

### Candidate UCOEs exhibit activity in a P19 silencing screen

We initially screened the candidates in the P19 murine embryonic carcinoma stem cell line. Murine embryonic carcinoma P19 cells are commonly used to study transgene silencing as they are susceptible to silencing within 2-3 weeks while other cell lines can take months. Early studies and characterization of A2UCOE in P19 cells support that it is a valid model system for studying anti-silencing activity that is predictive of efficacy in other cells and *in vivo* (14,36). As P19 cells readily integrate DNA, we performed a stable transfection of the expression construct. The EF1α (Elongation Factor 1) promoter was selected as the promoter to be linked with the candidate UCOEs because of its non-viral origin, so as to disregard the effect of viral recognition silencing (37), and its high expression level, which may allow for better dynamics in identifying the best performing candidates. Because stable transfections have a low efficiency of integration, a selection cassette was incorporated into the construct. We selected another endogenous non-viral promoter, the hPGK promoter, to drive expression of the puromycin resistance gene, and designed the EF1α-GFP and hPGK-PuroR cassettes to be oriented in opposing directions with the polyA terminators back-to-back, for maximal separation between the two promoters (Figure 2a). This design was intended to reduce polymerase run-through from one cassette to the other, as well as maintain genetic distance so that the epigenetic mechanism might be able to act independently on each promoter.

**Figure 2.**
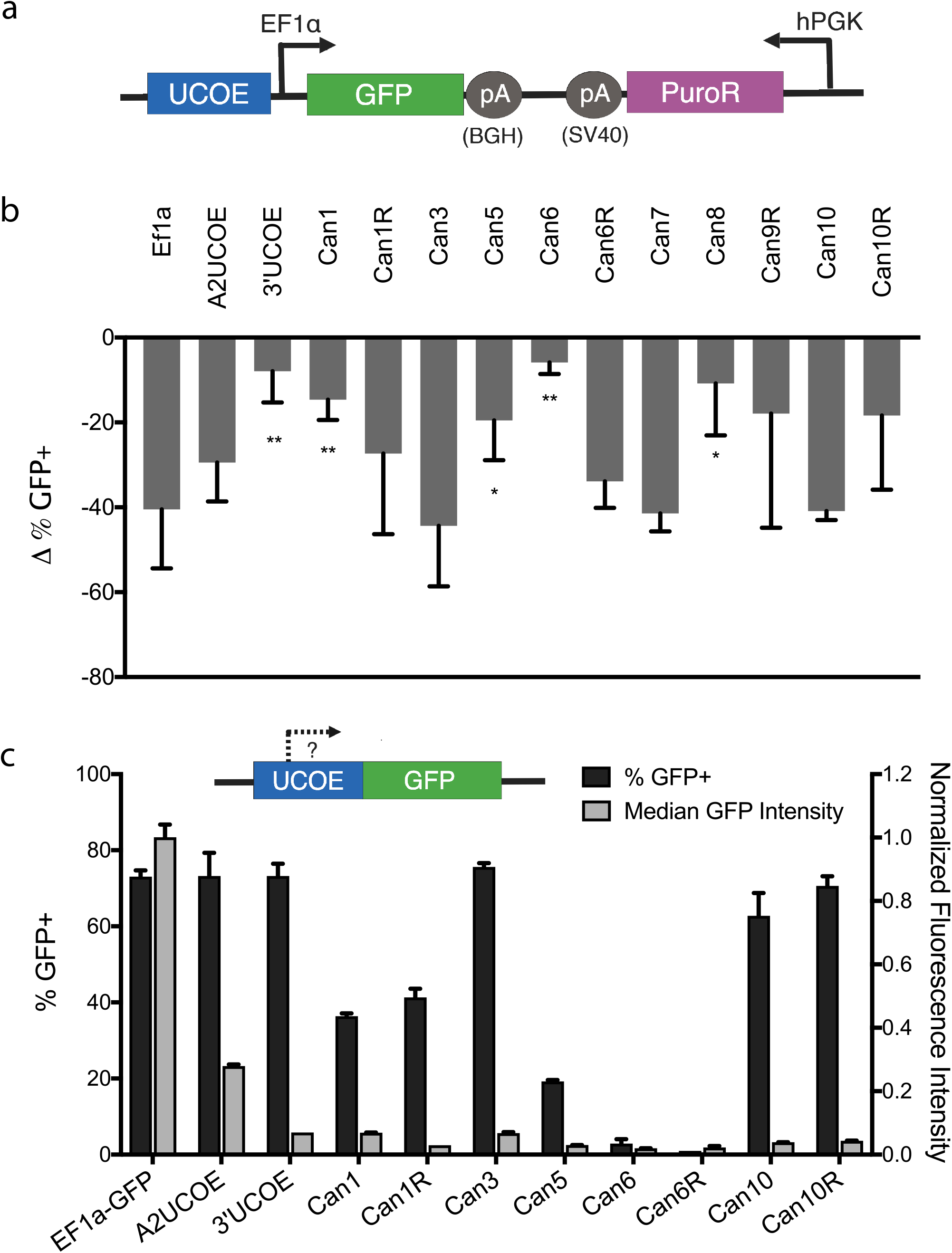
Screen of putative UCOEs in P19 embryonal carcinoma cells. a) Schematic of the dual expression construct used for screening putative UCOEs for anti-silencing activity in stable transfections. The GFP cassette is driven by the Elongation Factor 1 (EF1α) promoter and followed by the bovine growth hormone polyadenylation signal (BGH pA). The puromycin resistance cassette is oriented in the opposite direction, and driven by the human phosphoglycerate kinase (hPGK) promoter and followed by the Simian virus 40 polydenylation signal (SV40 pA). b) Silencing expression data of UCOE candidates linked to the EF1α promoter after stable transfection of P19 cell lines. After 2 weeks of selection for integrants, puromycin is removed to initiate silencing and cells are analyzed over a period of 19 days. Data shown is the difference in the percent GFP positive between the initial (day 0) timepoint and day 18. Data is reported as mean ± SD from at least three biological replicates. Asterisks indicate significance from an unpaired t-test with the no insulator control (* p<0.05, ** p< 0.005). c) Characterization of promoter activity of UCOE candidates. Stably transfected cells were assayed for % GFP+ and median fluorescent intensity (normalized to EF1α promoter control). Data is reported as mean ± SD from biological duplicates.

Candidate UCOEs were cloned directly upstream of the EF1α-EGFP cassette after PCR from a genome prep of the U2OS human osteosarcoma cell line. Candidates 2 and 4 were not recoverable with PCR. As a positive control, the 2.2 kb A2UCOE sequence, as well as the 1.2 kb reverse orientation sequence 3’UCOE (Figure 1b), were cloned into the same reporter construct as the candidate sequences. All candidate constructs and controls were transfected into P19 cells and selected for stable integrants by passaging in antibiotic-selective media over two weeks. After two weeks of selection, cells were transferred into antibiotic-free media to relieve the selection pressure that would counteract gene silencing. Cells were passaged every 2-3 days and the percent GFP positive in the population was monitored as a metric for silencing by flow cytometry analysis at each passage. The results demonstrate an exponential decay in this metric over the course of 19 days, with the negative (no-insulator) control having the most drastic decay over the first five days while active insulators resulted in more consistent gene expression over time (Supplementary Figure 2). Analysis of the loss of GFP-positive cells over time (Figure 2b) showed that both A2UCOE and 3’UCOE conferred silencing resistance compared to the negative control, with the 1.2 kb 3’UCOE mediating only an 8% loss in percent GFP positive cells over the time period, compared to 40% for the negative control and 29% for the 2.2 kb A2UCOE sequence. Additionally, four of the eleven tested candidates, Candidates 1, 5, 6 and 8, showed a significant improvement in stable expression relative to the negative control. In particular, Candidate 6 conferred the least loss of expression at about 6% loss over the 19 days, although the reverse orientation of Candidate 6 conferred no protective effect. Absolute expression, measured by GFP intensity, was not substantially different across cell populations harboring different controls and candidate UCOEs, confirming gene silencing as an all-or-nothing per-cell phenomenon (Supplementary Figure 3).

As A2UCOE dually functions as a protective regulatory element and a universal promoter, we further screened the candidate UCOEs for standalone promoter activity. We performed a similar experiment as the aforementioned screen, using a similar characterization construct that lacked the EF1α promoter (inlay, Figure 2C). This construct was used to assess whether the candidate sequences could drive reporter expression and act as stand-alone functional promoters. Candidates were compared to the positive control of the EF1α promoter, which had ~70% GFP+ cells after two weeks of antibiotic selection, showing a distinct positive population in the fluorescence histogram (Supplementary Figure 4). We consider a % GFP+ of at least 50% to reflect the existence of a second population in the histograms, and thus a binary indicator of promoter function. The median fluorescence intensity of the GFP+ population was used a measure of the strength of the promoter, and normalized to the EF1α positive control (Figure 2C). As expected, A2UCOE and 3’UCOE both exhibit promoter activity, with A2UCOE driving more than twice the absolute expression of GFP as 3’UCOE (Figure 2C). This aligns with previous observations that the *RNPA2B1* promoter is much stronger than that of *CBX3* (14). However, the *RNPA2B1* promoter in A2UCOE is only 30% as strong as the EF1α promoter. Several of the candidate UCOEs tested, including Candidates 3, 10 and 10R, are promoters comparable to 3’UCOE with absolute expression of about 10% of EF1α. On the other hand, Candidates 6 and 6R did not exhibit promoter activity, having expression at the level of background noise at 1-3% GFP+. Candidate 6 encompasses the entire regulatory region between the transcription start sites of the divergent genes *SURF1* and *SURF2*, as well as the first exon and intron of each gene (Figure 3). Therefore, this region must encompass the endogenous promoters for *SURF1* and *SURF2*. However, in our synthetic construct, the promoters are non-functional, suggesting that the promoter activity is dependent on an enhancer sequence that might be quite distant in two-dimensional sequence space but topologically close in three-dimensional space. Because of the nonmodularity and potential safety concerns associated with A2UCOE’s bidirectional promoter activity, Candidate 6 may present advantages in particular applications by not exhibiting promoter activity.

**Figure 3.**
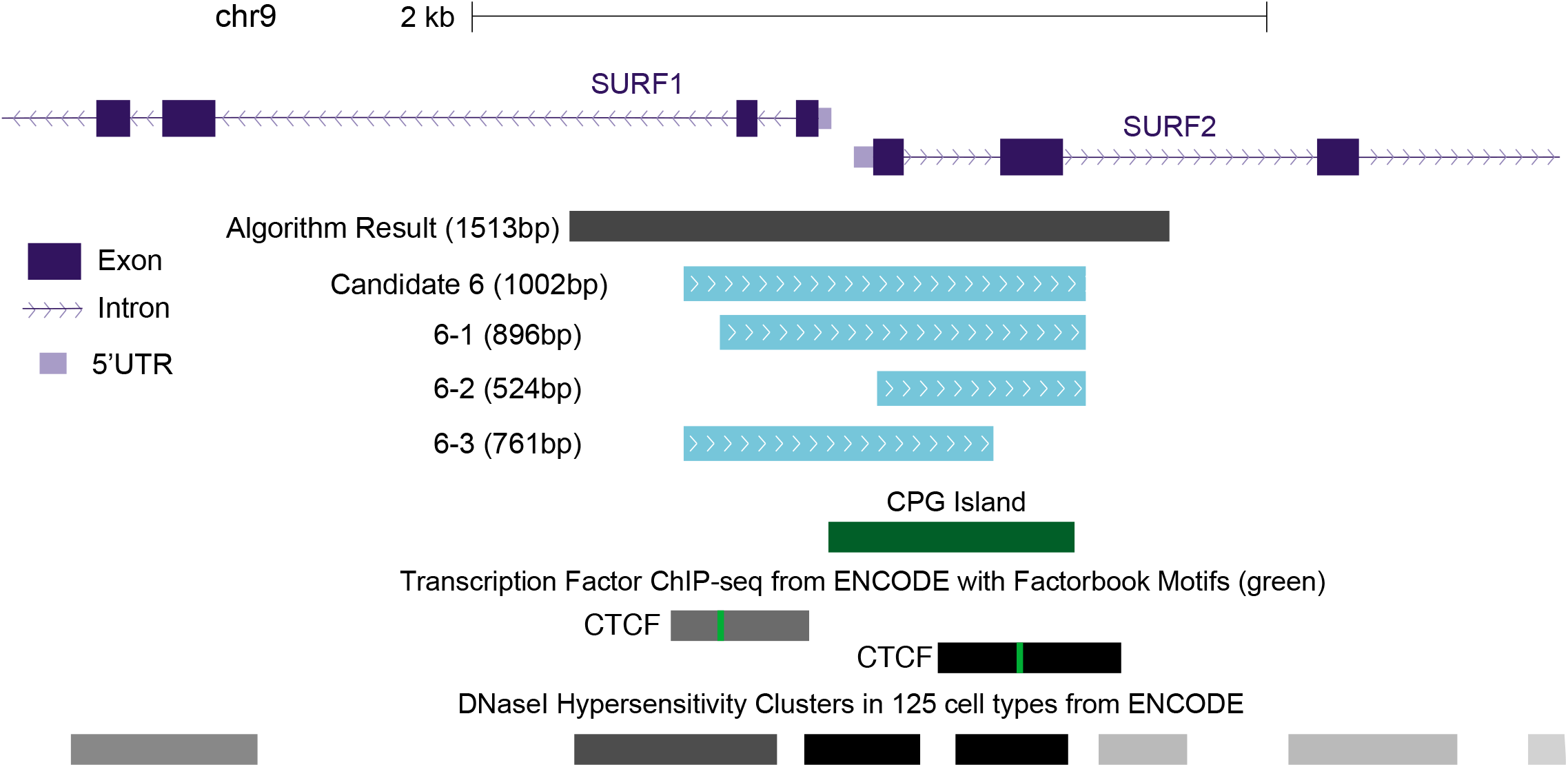
Illustration of Candidate 6 locus (human SURF1/SURF2 genomic region) and associated truncations. Algorithm result for Candidate 6 points to the locus encompassing the start of the *SURF1* and *SURF2* divergently oriented genes, shown in this adapted screenshot from the UCSC Genome Browser. Boundaries for the 1,002 bp Candidate 6 were drawn to include the entirety of the CpG island and CTCF sites. Truncation 6-1 shortens the 5’ end but maintains all identified features of the locus. Truncation 6-2 further truncates the 5’ end such that it lacks the first CTCF binding site and excludes the intergenic region between SURF1 and SURF2. Truncation 6-3 is truncated from the 3’ end to exclude the second CTCF site.

### Candidate 6 and associated truncations demonstrate activity across multiple promoters

While convenient for a screen, the stable transfection methodology is uncontrolled for copy number, and integration sites are likely biased by antibiotic selection. Thus, we chose a more reproducible and applicable integration technology in lentiviral transduction to further characterize the most active candidate UCOE element Candidate 6. We constructed a series of lentiviral constructs that paired the candidate UCOE regions with four commonly used mammalian promoters (Figure 4a). Candidate 6 spans the entire region between *SURF1* and *SURF2* including the first introns of both genes. In an effort to identify the core functional region of Candidate 6 and determine shorter sequences that exhibit this activity, we further examined three truncated versions of Candidate 6 in this assay (Figure 3). Truncation 6-1 was designed to keep as many of the important features as possible while removing additional spacer sequence; thus, the 5’ end of the Candidate 6 region was truncated up to the first CTCF binding site and the CpG island kept completely intact. Truncation 6-2 was designed to incorporate a larger truncation of the 5’ end of the element to remove the intergenic region between *SURF1* and *SURF2* and the first CTCF binding site while still incorporating the majority of the CpG island. Truncation 6-3 was truncated at the 3’ end of the element, thereby removing the second CTCF binding site. We hypothesized that 6-1 would function as well as Candidate 6 due to the retention of most of the hypothesized functional features. We expected that 6-3 would exhibit reduced efficacy due to its loss of a CTCF binding site, and were particularly interested in whether 6-2, which lacked the intergenic regulatory area between the divergent *SURF1* and *SURF2* genes, would maintain anti-silencing activity.

**Figure 4.**
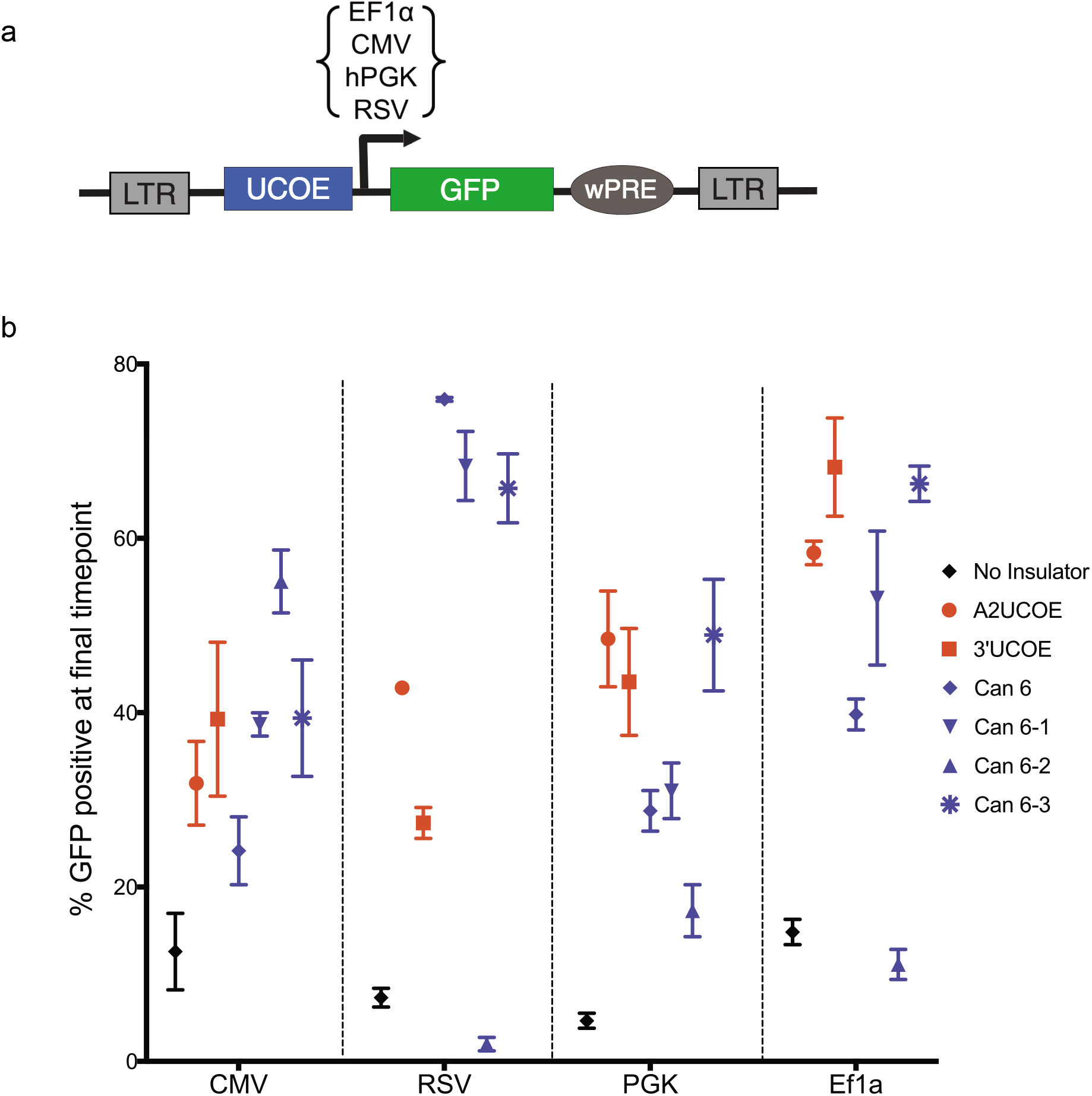
Candidate 6 resists transgene silencing from lentiviral transductions across four different promoters. (a) Schematic of the lentiviral donor construct for assaying putative UCOE Candidate 6 and truncations for anti-silencing activity in stable transductions. The lentiviral expression cassette includes the wPRE posttranscriptional regulatory element, and is flanked by two long terminal repeat (LTR) sequences. (b) Percent GFP positive cells at the final timepoint (day 26 for CMV and RSV, day 27 for PGK and EF1α) of the lentivirus silencing experiment across the four tested promoters. P19 cells were FACS-sorted five days after transduction (day 0), and then assayed over a 26 or 27-day time period thereafter. Data is reported as mean ± SD from three biological replicates. Representative SSC-A vs GFP plots are shown in Supplementary Figure 6.

P19 cells were transduced with lentiviral constructs harboring the four Candidate 6 regions, two positive UCOE controls (A2UCOE, 3’UCOE), and a negative control (no insulator region). Transduced cells were FACS-sorted after lentiviral integration at a low MOI to ensure single integrants (MOIs in Supplementary Table 3), and the percent GFP positive cells were assayed over time as with the stable transfection experiments, demonstrating exponential decay over time (Supplementary Figure 5). As all conditions were FACS-sorted at day 0 to 100% GFP positive, the percent GFP positive cells at day 26 (CMV/RSV) or day 27 (PGK/ EF1α) is a measurement of the amount of silencing that has occurred (Figure 4b, representative flow plots in Supplementary Figure 6). When linked to EF1α, A2UCOE, 3’UCOE, and Candidate 6-3 have more than 4 times the GFP positive cells at day 26 compared to the negative control, with greater than 65% of cells GFP positive at d27 for Candidate 6-3 compared to 15% in the negative control. The full-length Candidate 6 and 6-1 element maintain 2.7 and 3.6 times the expression of the negative control, respectively. Meanwhile, 6-2 is ineffective, at 11% GFP+ cells compared to 15% in the negative control.

For the CMV promoter, the negative control shows about 13% GFP+ cells at day 26. Unlike the other promoters tested, Candidate 6-2 is the best-performing population in this promoter, mediating 55% GFP+, a 4.4-fold improvement over the negative control and a 1.4-fold improvement over A2UCOE. A2UCOE, 6-1, and 6-3, all perform equivalently with 39% GFP+ cells at day 26, which is a 3-fold improvement over the negative control. Here, the full-length A2UCOE is slightly more effective than 3’UCOE, and is exactly matched by truncations 6-1 and 6-3 at 39% GFP+.

For the PGK promoter, only about 5% of the negative control cells are still GFP+ at day 27. A2UCOE, 3’UCOE, and Candidate 6-3 perform similarly, maintaining a %GFP+ 8-fold higher than the control at day 27, with Candidate 6-3 mediating 49% GFP positive cells at the final timepoint. The full-length Candidate 6 and truncation 6-1 demonstrate 29% and 31% GFP+ at day 27, respectively, which corresponds to a 5-fold improvement over the negative control. Candidate 6-2 is substantially less effective than the other Candidate 6 sequences at 17% GFP+ cells at day 27.

Finally, Candidate 6 and associated truncations demonstrate the most improvement over the A2UCOE elements in the RSV promoter construct. At day 26, only 7% of cells in the negative control remain GFP+. A2UCOE and 3’UCOE exhibit substantial improvement over the control at 43% and 27% GFP+, respectively. Markedly, Candidate 6 maintains 76% GFP+ cells, with truncations 6-1 and 6-3 exhibiting 68% and 66% GFP+ cells, respectively. These three elements show at least a 9-fold improvement over the negative control and at least 1.5-fold over A2UCOE and 2.4-fold over the 1.2 kb 3’UCOE. Truncation 6-2, on the other hand, is ineffective when linked to the RSV promoter.

Taken together, the data demonstrate that Candidate 6 and its associated truncations (with the exception of truncation 6-2) showed substantial improvement over the negative control across all four tested promoters and performed on par (PGK/ EF1α) or at least 1.4 times better (CMV/RSV) than the positive controls A2UCOE and 3’UCOE. Candidate 6 and associated truncations were most efficacious in concert with the RSV promoter, outperforming the 2.2 kb A2UCOE by 1.5-fold, and the 1.2 kb 3’UCOE by more than two-fold in percent GFP+ cells after 26 days. While there is variability in the performance of the Candidate 6 truncations depending on the promoter, Candidate 6-3 exhibits the most consistent activity, outperforming the full-length Candidate 6 sequence in all promoters except RSV (where it is still highly effective). Thus, we suggest that the 767 bp Candidate 6-3 element would be an effective first choice for researchers looking to mediate anti-silencing activity, as this element maintains at least an equivalent level of percent GFP+ cells as A2UCOE/3’UCOE across all four promoters tested. Notably, truncation 6-2, which completely lacks the intergenic area between *SURF1* and *SURF2* genes, failed to outperform the negative control in 3 of the 4 promoters tested, suggesting that the functional core of the element is located within this intergenic region. The notable exception to this is the substantial protective effect of the 6-2 element with the CMV promoter, which indicates that the particular interplay of the 6-2 sequence and the components of the CMV promoter combine for a unique protective effect.

### Effective UCOEs confer resistance to DNA CpG methylation and histone deacetylation

We next examined whether Candidate 6 functioned on an epigenetic level to resist transgene silencing. It is well understood that transgene silencing is mediated by the loss of histone acetylation at the locus and addition of DNA methylation (4). Two small molecule drugs have been widely used to probe this effect, trichostatin A (TSA), which is a specific inhibitor of histone deacetylase, and 5-azacytidine (5-aza), a cytidine analog that inhibits methylation upon its incorporation into DNA. Both molecules have been individually used to reactivate expression of silenced transduced genes and to conclude that histone deacetylation and CpG methylation are integrally involved in transgene silencing (38–40).

Transduced P19 cells undergoing the previously described silencing experiment were replica plated at late passages and treated a day later with a range of concentrations of 5-aza or TSA. Twenty-four hours later, cells were assayed by flow cytometry for reactivation of GFP expression (Figure 5, Supplementary Figure 7). All examined populations showed a dose-dependent increase in %GFP positive cells, with the highest dose of 5-aza rescuing 63% of the silenced cells in the RSV promoter-only construct. For every condition tested, increasing concentration of 5-aza increased the fraction of silenced cells that were reactivated, suggesting that even more cells may have been susceptible to 5-aza rescue if the toxicity of the chemical had not limited the concentration. This effect corroborates that the silencing seen in transduced cells is due to methylation of the DNA construct. Similarly, TSA-treated cells show a dose-dependent recovery of GFP expression, although to a smaller extent than 5-aza, with the highest dose of TSA rescuing only about 25% of silenced cells in the RSV promoter-only construct. These results further confirm the role of histone deacetylation in the silencing of transduced P19 cells. Even across the most effective UCOEs (in RSV, Candidate 6), more than 80% of silenced cells at day 18 and later can be rescued with the small molecule effectors. Taken together, the data indicate that silencing in our transduction experiments is due to epigenetic effects as opposed to a loss of the DNA construct, and that UCOEs (and particularly Candidate 6 and truncations) function by resisting DNA methylation or histone deacetylation at the integration locus.

**Figure 5.**
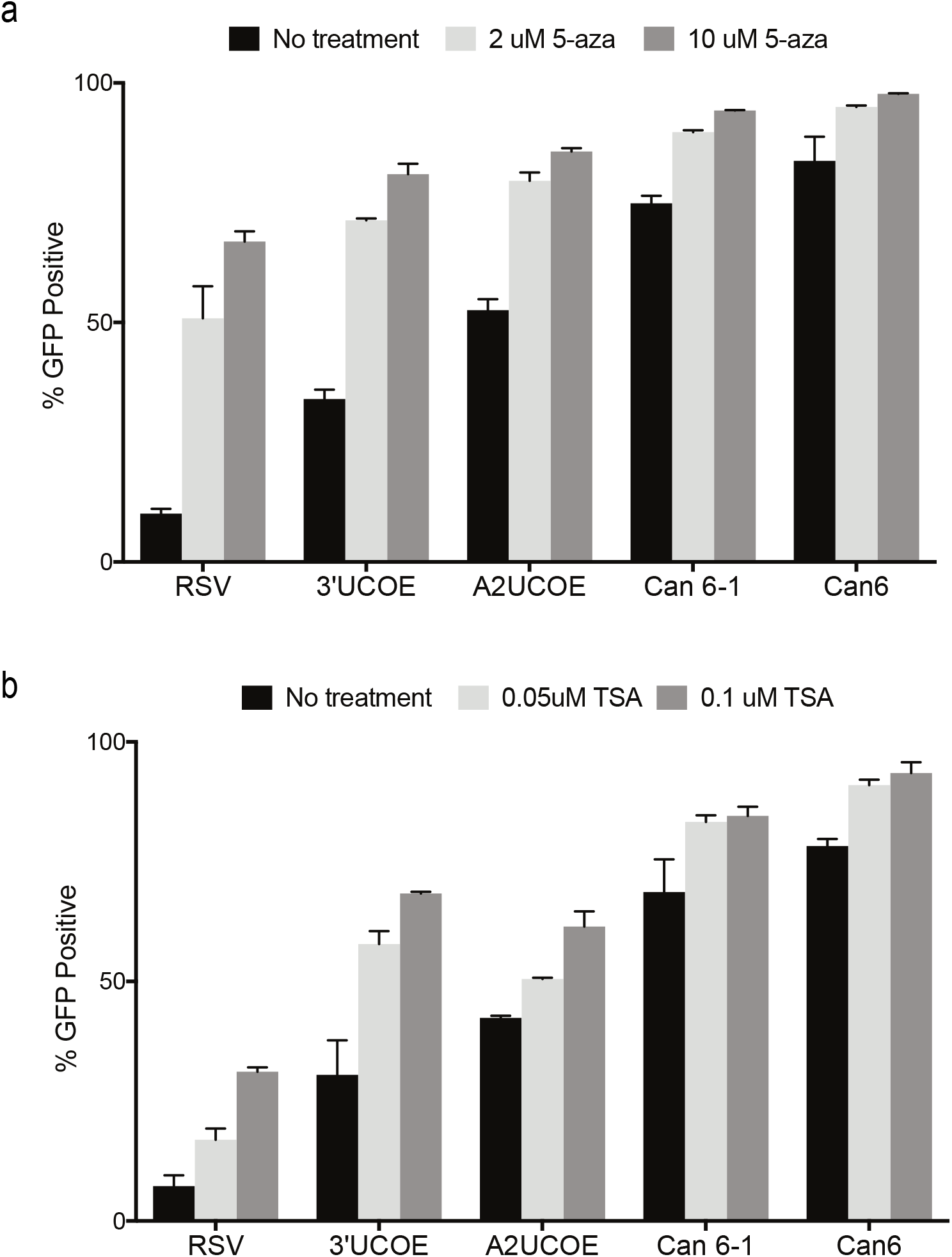
Candidate 6 resists DNA methylation and histone deacetylation. (a) GFP expression is rescued by treatment with DNA methylation inhibitor 5-aza-cytidine (5-aza) in day 18 UCOE-RSV cells from the lentiviral silencing experiment. Cells were replica plated at day 16, specified concentrations of 5-aza were introduced 24 hours later (with exception of control), and cells were passaged and assayed via flow cytometry 24 hours after chemical introduction for % GFP+ cells. Data is reported as mean ± SD from three biological replicates. (b) GFP expression is rescued by treatment with HDAC inhibitor trichostatin A (TSA) in day 24 UCOE-RSV cells from the lentiviral silencing experiment. Cells were replica plated at day 22, specified concentrations of TSA were introduced 24 hours later (with exception of control), and cells were passaged and assayed via flow cytometry 24 hours after chemical introduction for % GFP+ cells. Data is reported as mean ± SD from three biological replicates.

## DISCUSSION

The eukaryotic genome is a highly complex, interconnected epigenetic environment that subjects transgenes to time-dependent epigenetic silencing. A2UCOE and its associated truncations have been demonstrated to prevent silencing and mediate stable expression across many applications. Here, we demonstrate the discovery of novel putative UCOEs from a feature-driven search based on the natural locus of A2UCOE. From our computational search of the human genome, we discovered an element from the SURF housekeeping gene locus (Candidate 6) that we name SRF-UCOE. In transduced P19 cells, SRF-UCOE prevents silencing in at least 1.4-fold more cells than the previously characterized A2UCOE when linked with RSV or CMV promoters, and matches the performance of A2UCOE when paired with the EF1α or PGK promoters, mediating at least a 4fold improvement in GFP positive cells compared to the promoter-only control. Although the 761 bp 6-3 truncation lacks the second CTCF binding site present in the full-length 1 kb SRF-UCOE sequence, we observed that it stabilized expression in a higher percentage of cells than the full-length sequence in 3 of the 4 promoters examined in this work. Further study is needed to better understand why the exclusion of the second exon of *SURF2* (and its corresponding CTCF site) would improve the UCOE’s functionality.

We observed wide variability in UCOE protective effects depending on the promoter sequence used. For example, when pairing any of the UCOEs, including A2UCOE, with the PGK promoter, we were not able to maintain more than 50% GFP positive cells after 27 days of study. In contrast, the SRF-UCOE was able to maintain more than 70% GFP positive cells after 26 days of study when paired with the RSV promoter. The observed variability in effect across promoters for both A2UCOE and SRF-UCOE suggests that the mechanism of silencing is slightly different for different promoters, such that the same UCOE does not exhibit a consistent effect across these different contexts, and highlights the importance of having an expanded collection of functional UCOEs.

The human Surfeit housekeeping locus is a unique, highly conserved cluster of six housekeeping genes. The orientation of each gene alternates from its neighbor, making it a locus of multiple divergent housekeeping gene promoters, which has been considered an important hallmark of UCOEs (41). Most notably, the locus is unique because of the extremely short distances between the genes in the cluster (between 97 and 374 bp), making it one of the tightest known mammalian gene clusters (42). These features support the identification of a strong chromatin remodelling element at this locus. We speculate that further study of the Surfeit locus, including the *SURF3-SURF4* intergenic region, may identify additional UCOE elements.

The SRF-UCOE region tested here spans the first introns of *SURF1* and *SURF2* and the entirety of the genomic sequence between them, implying the inclusion of an endogenous bidirectional promoter. However, the Candidate 6 region does not initiate transcription in the synthetic constructs tested in this study. These results suggest that the endogenous promoter activity of this region must be mediated by additional environmental components. As with most housekeeping genes, this region does not contain a TATA box, but has a strongly predicted SP2 transcription factor binding site according to the comprehensive database of TFBS binding profiles, JASPAR (43). Future experiments may be focused on elucidating the mechanism of its endogenous but non-universal transcriptional profile. The lack of inherent transcriptional activity in non-natural constructs makes the element a more modular part when paired with promoters with desired strengths for a given application, as well as reducing the possibility of unwanted off-target effects of a bidirectional promoter upon random integration, a previously identified disadvantage of A2UCOE (16). Our results contrast with previous hypotheses that the mechanism of chromatin opening for UCOEs is directly tied to its mediation of bidirectional transcriptional activity (13,17). Our results suggest that bidirectional promoters can be an indicator for UCOEs, but are not the driving mechanism for their chromatin remodelling activity. Previous studies that have truncated 3’UCOE to contain merely the *CBX3* promoter while maintaining anti-silencing properties (16) further support the conclusion that bidirectional transcriptional activity is not necessary for the observed effect on silencing, but here we demonstrate the first example to-date of a UCOE without any inherent transcriptional activity. This finding reiterates how the discovery of more UCOEs can improve our understanding of their mechanism of action.

In this work, we demonstrated the utility of a feature-driven algorithm for discovering novel genetic parts that we expect could be generalized to discovering other natural genetic parts such as promoters and enhancers. Using this approach, we identified and characterized a novel UCOE (SRF-UCOE) that is the first example to-date of a UCOE without inherent transcriptional activity. We expect this unique feature of SRF-UCOE to expand its utility for clinically relevant gene and cell therapy applications where off-target effects are a concern. Additionally, in this study, we show the effectiveness of a core 761 bp sequence of this UCOE, and we believe the shorter sequence can be enabling for retroviral and adenoviral vectors in gene therapy where there are critical size limitations.

In our hands, the 1 kb SRF-UCOE resists silencing in at least 1.5 times the number of cells than the previously-available 2.2 kb A2UCOE, and in 2.4 times the number of cells than the 1.2 kb 3’UCOE, when paired with the RSV promoter. We predict that as this new element continues to be tested, there will likely be other specific promoters and cell types where SRF-UCOE can outperform other available UCOEs. In this way, the expanding toolbox of UCOEs will pave the way for expanded use of these extremely useful tools for mediating stable long-term expression. Finally, as synthetic biology circuits come of age in cell therapies and molecular diagnostics, UCOEs will make possible the use of cellular logic over long-term applications.

## DATA AVAILABILITY

Linux shell scripts for the computational algorithm are available on Github (www.github.com/srudina/UCOEdiscovery).

## SUPPLEMENTARY DATA

Supplementary Data are available at NAR online.

## ACKNOWLEDGEMENT

We thank B. Kotopka, J. Xiang, A. Cravens, C. Kim, and P. Dykstra for valuable comments on the manuscript. Cell sorting for this project was performed on instruments in the Stanford Shared FACS Facility, with significant technical assistance from M. Weglarz. We thank T. Bauer, G. Bejerano and J. Notwell for their advice and input in the computational component of this project. We thank Dr. Fang Zhang of University College, London for the gift of A2UCOE plasmid.

## REFERENCES

1. Jostock, T. and Knopf, H.P. (2012) Mammalian stable expression of biotherapeutics. Methods Mol Biol, 899, 227–238.

2. Yang, Y., Mariati, Chusainow, J. and Yap, M.G. (2010) DNA methylation contributes to loss in productivity of monoclonal antibody-producing CHO cell lines. J Biotechnol, 147, 180–185.

3. Oleg E. Tolmachov, T.S.a.T.T. (2013) In Molina, F. M. (ed.), Gene Therapy. IntechOpen.

4. Alhaji, S.Y., Ngai, S.C. and Abdullah, S. (2018) Silencing of transgene expression in mammalian cells by DNA methylation and histone modifications in gene therapy perspective. Biotechnol Genet Eng Rev, 1–25.

5. Ellis, J. (2005) Silencing and variegation of gammaretrovirus and lentivirus vectors. Human Gene Therapy, 16, 1241–1246.

6. Emery, D.W. (2011) The use of chromatin insulators to improve the expression and safety of integrating gene transfer vectors. Hum Gene Ther, 22, 761–774.

7. Chung, J.H., Bell, A.C. and Felsenfeld, G. (1997) Characterization of the chicken beta-globin insulator. Proc Natl Acad Sci U S A, 94, 575–580.

8. Arumugam, P.I., Scholes, J., Perelman, N., Xia, P., Yee, J.K. and Malik, P. (2007) Improved human beta-globin expression from self-inactivating lentiviral vectors carrying the chicken hypersensitive site-4 (cHS4) insulator element. Mol Ther, 15, 1863–1871.

9. Arumugam, P.I., Urbinati, F., Velu, C.S., Higashimoto, T., Grimes, H.L. and Malik, P. (2009) The 3’ region of the chicken hypersensitive site-4 insulator has properties similar to its core and is required for full insulator activity. PLoS One, 4, e6995.

10. Otte, A.P., Kwaks, T.H., van Blokland, R.J., Sewalt, R.G., Verhees, J., Klaren, V.N., Siersma, T.K., Korse, H.W., Teunissen, N.C., Botschuijver, S. et al. (2007) Various expression-augmenting DNA elements benefit from STAR-Select, a novel high stringency selection system for protein expression. Biotechnol Prog, 23, 801–807.

11. Izumi, M. and Gilbert, D.M. (1999) Homogeneous tetracycline-regulatable gene expression in mammalian fibroblasts. J Cell Biochem, 76, 280–289.

12. Saunders, F., Sweeney, B., Antoniou, M.N., Stephens, P. and Cain, K. (2015) Chromatin function modifying elements in an industrial antibody production platform--comparison of UCOE, MAR, STAR and cHS4 elements. PLoS One, 10, e0120096.

13. Antoniou, M., Harland, L., Mustoe, T., Williams, S., Holdstock, J., Yague, E., Mulcahy, T., Griffiths, M., Edwards, S., Ioannou, P.A. et al. (2003) Transgenes encompassing dual-promoter CpG islands from the human TBP and HNRPA2B1 loci are resistant to heterochromatin-mediated silencing. Genomics, 82, 269–279.

14. Zhang, F., Frost, A.R., Blundell, M.P., Bales, O., Antoniou, M.N. and Thrasher, A.J. (2010) A ubiquitous chromatin opening element (UCOE) confers resistance to DNA methylation-mediated silencing of lentiviral vectors. Mol Ther, 18, 1640–1649.

15. Dighe, N., Khoury, M., Mattar, C., Chong, M., Choolani, M., Chen, J., Antoniou, M.N. and Chan, J.K. (2014) Long-term reproducible expression in human fetal liver hematopoietic stem cells with a UCOE-based lentiviral vector. PLoS One, 9, e104805.

16. Muller-Kuller, U., Ackermann, M., Kolodziej, S., Brendel, C., Fritsch, J., Lachmann, N., Kunkel, H., Lausen, J., Schambach, A., Moritz, T. et al. (2015) A minimal ubiquitous chromatin opening element (UCOE) effectively prevents silencing of juxtaposed heterologous promoters by epigenetic remodeling in multipotent and pluripotent stem cells. Nucleic Acids Res, 43, 1577–1592.

17. Brendel, C., Muller-Kuller, U., Schultze-Strasser, S., Stein, S., Chen-Wichmann, L., Krattenmacher, A., Kunkel, H., Dillmann, A., Antoniou, M.N. and Grez, M. (2012) Physiological regulation of transgene expression by a lentiviral vector containing the A2UCOE linked to a myeloid promoter. Gene Ther, 19, 1018–1029.

18. Haenseler, W., Kuzmenko, E., Smalls-Mantey, A., Browne, C., Seger, R., James, W., & Siler, U. (2018) Lentiviral gene therapy vector with UCOE stably restores function in iPSC-derived neutrophils of a CDG patient. Matters.

19. Benton, T., Chen, T., McEntee, M., Fox, B., King, D., Crombie, R., Thomas, T.C. and Bebbington, C. (2002) The use of UCOE vectors in combination with a preadapted serum free, suspension cell line allows for rapid production of large quantities of protein. Cytotechnology, 38, 43–46.

20. Williams, S., Mustoe, T., Mulcahy, T., Griffiths, M., Simpson, D., Antoniou, M., Irvine, A., Mountain, A. and Crombie, R. (2005) CpG-island fragments from the HNRPA2B1/CBX3 genomic locus reduce silencing and enhance transgene expression from the hCMV promoter/enhancer in mammalian cells. BMC Biotechnol, 5, 17.

21. Adamson, B., Norman, T.M., Jost, M., Cho, M.Y., Nunez, J.K., Chen, Y., Villalta, J.E., Gilbert, L.A., Horlbeck, M.A., Hein, M.Y. et al. (2016) A Multiplexed Single-Cell CRISPR Screening Platform Enables Systematic Dissection of the Unfolded Protein Response. Cell, 167, 1867–1882 e1821.

22. Jost, M., Chen, Y., Gilbert, L.A., Horlbeck, M.A., Krenning, L., Menchon, G., Rai, A., Cho, M.Y., Stern, J.J., Prota, A.E. et al. (2017) Combined CRISPRi/a-Based Chemical Genetic Screens Reveal that Rigosertib Is a Microtubule-Destabilizing Agent. Mol Cell, 68, 210–223 e216.

23. Uchiyama, T., Adriani, M., Jagadeesh, G.J., Paine, A. and Candotti, F. (2012) Foamy virus vector-mediated gene correction of a mouse model of Wiskott-Aldrich syndrome. Mol Ther, 20, 1270–1279.

24. Kunkiel, J., Godecke, N., Ackermann, M., Hoffmann, D., Schambach, A., Lachmann, N., Wirth, D. and Moritz, T. (2017) The CpG-sites of the CBX3 ubiquitous chromatin opening element are critical structural determinants for the anti-silencing function. Sci Rep, 7, 7919.

25. Yi, Y., Noh, M.J. and Lee, K.H. (2011) Current advances in retroviral gene therapy. Curr Gene Ther, 11, 218–228.

26. Ernst, J., Kheradpour, P., Mikkelsen, T.S., Shoresh, N., Ward, L.D., Epstein, C.B., Zhang, X., Wang, L., Issner, R., Coyne, M. et al. (2011) Mapping and analysis of chromatin state dynamics in nine human cell types. Nature, 473, 43–49.

27. Hsu, F., Kent, W.J., Clawson, H., Kuhn, R.M., Diekhans, M. and Haussler, D. (2006) The UCSC Known Genes. Bioinformatics, 22, 1036–1046.

28. Consortium, E.P. (2012) An integrated encyclopedia of DNA elements in the human genome. Nature, 489, 57–74.

29. Wang, J., Zhuang, J., Iyer, S., Lin, X.Y., Greven, M.C., Kim, B.H., Moore, J., Pierce, B.G., Dong, X., Virgil, D. et al. (2013) Factorbook.org: a Wiki-based database for transcription factor-binding data generated by the ENCODE consortium. Nucleic Acids Res, 41, D171–176.

30. She, X., Rohl, C.A., Castle, J.C., Kulkarni, A.V., Johnson, J.M. and Chen, R. (2009) Definition, conservation and epigenetics of housekeeping and tissue-enriched genes. BMC Genomics, 10, 269.

31. Kent, W.J., Sugnet, C.W., Furey, T.S., Roskin, K.M., Pringle, T.H., Zahler, A.M. and Haussler, D. (2002) The human genome browser at UCSC. Genome Research, 12, 996–1006.

32. Zufferey, R. and Trono, D. (2000) Production of High-Titer Lentiviral Vectors. Current Protocols in Human Genetics, 26, 12.10.11–12.10.12.

33. Chen, S., Sanjana, N.E., Zheng, K., Shalem, O., Lee, K., Shi, X., Scott, D.A., Song, J., Pan, J.Q., Weissleder, R. et al. (2015) Genome-wide CRISPR screen in a mouse model of tumor growth and metastasis. Cell, 160, 1246–1260.

34. Lindahl Allen, M. and Antoniou, M. (2007) Correlation of DNA methylation with histone modifications across the HNRPA2B1-CBX3 ubiquitously-acting chromatin open element (UCOE). Epigenetics, 2, 227–236.

35. Weth, O., Paprotka, C., Gunther, K., Schulte, A., Baierl, M., Leers, J., Galjart, N. and Renkawitz, R. (2014) CTCF induces histone variant incorporation, erases the H3K27me3 histone mark and opens chromatin. Nucleic Acids Research, 42, 11941–11951.

36. Bannister, A.J. and Kouzarides, T. (2011) Regulation of chromatin by histone modifications. Cell Res, 21, 381–395.

37. Gill, D.R., Smyth, S.E., Goddard, C.A., Pringle, I.A., Higgins, C.F., Colledge, W.H. and Hyde, S.C. (2001) Increased persistence of lung gene expression using plasmids containing the ubiquitin C or elongation factor 1 alpha promoter. Gene Therapy, 8, 1539–1546.

38. Chen, W.Y., Bailey, E.C., McCune, S.L., Dong, J.Y. and Townes, T.M. (1997) Reactivation of silenced, virally transduced genes by inhibitors of histone deacetylase. Proc Natl Acad Sci U S A, 94, 5798–5803.

39. Pikaart, M.J., Recillas-Targa, F. and Felsenfeld, G. (1998) Loss of transcriptional activity of a transgene is accompanied by DNA methylation and histone deacetylation and is prevented by insulators. Genes Dev, 12, 2852–2862.

40. Kuriyama, S., Sakamoto, T., Kikukawa, M., Nakatani, T., Toyokawa, Y., Tsujinoue, H., Ikenaka, K., Fukui, H. and Tsujii, T. (1998) Expression of a retrovirally transduced gene under control of an internal housekeeping gene promoter does not persist due to methylation and is restored partially by 5-azacytidine treatment. Gene Ther, 5, 1299–1305.

41. Neville, J.J., Orlando, J., Mann, K., McCloskey, B. and Antoniou, M.N. (2017) Ubiquitous Chromatin-opening Elements (UCOEs): Applications in biomanufacturing and gene therapy. Biotechnol Adv, 35, 557–564.

42. Duhig, T. (1998) The human Surfeit locus. Genomics, 52, 72–78.

43. Khan, A., Fornes, O., Stigliani, A., Gheorghe, M., Castro-Mondragon, J.A., van der Lee, R., Bessy, A., Cheneby, J., Kulkarni, S.R., Tan, G. et al. (2018) JASPAR 2018: update of the open-access database of transcription factor binding profiles and its web framework. Nucleic Acids Res, 46, D1284.

