## Supplementary Information for "A Novel Chromatin-Opening Element for Stable Long-term Transgene Expression"

**Supplementary Text: Bioinformatics Algorithm Detailed Methods**

We downloaded the Broad peak data from ENCODE for GM12878, which spanned 13 histone modifications and transcription factors (Figure 1a). The files contain discrete intervals of ChIP-seq fragment enrichment through a statistical approach further described through the UCSC ENCODE portal, specifically using the Scripture software to call peaks (1), followed by a Matlab script to decouple smaller enriched intervals within very large intervals from the Scripture output. We used the bedtools intersect function consecutively on each of the 11 ChIP-seq signals that were associated with the A2UCOE locus: H3K4m1, H3K4m2, H3K4m3, H3K9ac, H3K27ac, H3K36m3, H3K79m2, H4K20m1. ChIP-seq peaks for EZH2, H2A.Z, and CTCF were also used in this fashion. We used bedtools subtract to remove any intervals in the resulting dataset for H3K9m3 and H3K27m3.

The UCSC Known Genes Track was downloaded from the UCSC genome browser. This Known Genes dataset was constructed based on protein data from Swiss-Prot and associated mRNA data from Genbank (2). We subtracted the known gene track from the working candidate list (using bedtools subtract) and then intersected that result with our working list (keeping the entirety of the original interval using bedtools intersect with the '-wa' option) to remove any regions that were completely within the known gene track.

The CpG island track was downloaded from the UCSC Genome Browser, which was generated using a modified version of a program developed by G. Miklem and L. Hillier, and predicts CpG islands using three particular criteria: (1) GC content of 50% or greater, (2) length greater than 200 bp, (3) ratio greater than 0.6 of observed number of CG dinucleotide to the expected number on the basis of the number of Gs and Cs in the segment. The program examines each base one at a time, scoring dinucleotides +17 for CG and –1 otherwise. This was intersected with the working list using bedtools intersect to find overlaps, and keep the entire original entry where there was a minimal overlap of 0.5 (50%).

DNA methylation data from Reduced Representation Bisulfite Sequencing (RRBS) in GM12878 was downloaded from ENCODE (3). This data consisted of intervals identified through RBSS, read counts within each interval, and the percent methylated CGs for each interval. We filtered this list first to only include entries with at least 10 reads, and then percent methylated (a) greater than 10%, (b) greater than 20%, and (c) less than 20%. We first used the overlapSelect function from UCSC to keep all candidates that overlapped with (a), but this criterion was too harsh, as it removed the A2UCOE locus from the list. Therefore, we used the overlapSelect function from UCSC to keep all candidates that overlapped with (c), and remove any that overlapped with (b). Finally, a simple overlapSelect was performed with the CTCF binding site data from ENCODE Transcription factor CHIP dataset, which encompasses data across several cell lines (4).

*Re-ranking candidate list by housekeeping gene coefficient of variance*

A list of identified housekeeping genes and their coefficients of variance was obtained from a 2009 study of the gene expression profiles of 42 tissues (5). The accession numbers were mapped to RefSeq gene names and their corresponding chromosome number and position. Using this file as an input along with our candidate list to the 'bedtools closest' function, we were able to obtain the distance to and identity of the closest housekeeping gene for each candidate, including regions that overlapped with the housekeeping gene. Results were then merged back with the coefficient of variations from the housekeeping-identifying study (5). The 88 regions were then sorted by distance to housekeeping gene, and then further sorted by the coefficients of variance to rank-order the candidate list. This ranking resulted in A2UCOE being at the top of the list (zeroth position), and candidates were named Candidate 1, 2, and onwards to result in a total of 87 putative UCOE candidates.

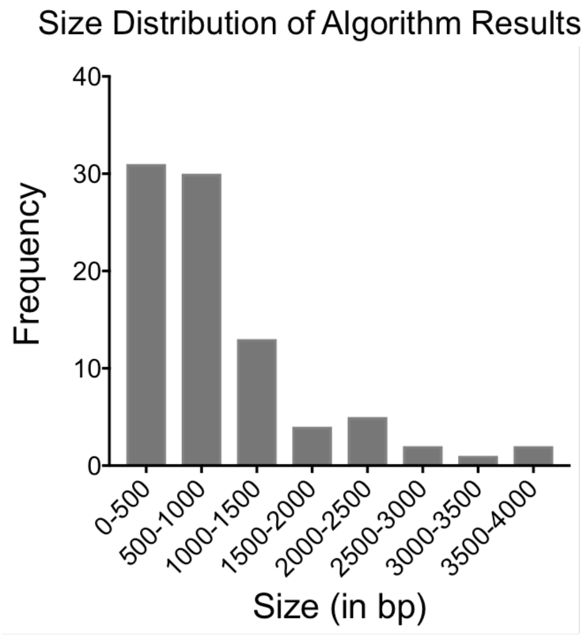

**Supplementary Figure 1**. **Size distribution of the 87 candidate UCOE regions returned by the algorithm.** The majority of regions identified were under 1,000 bp.

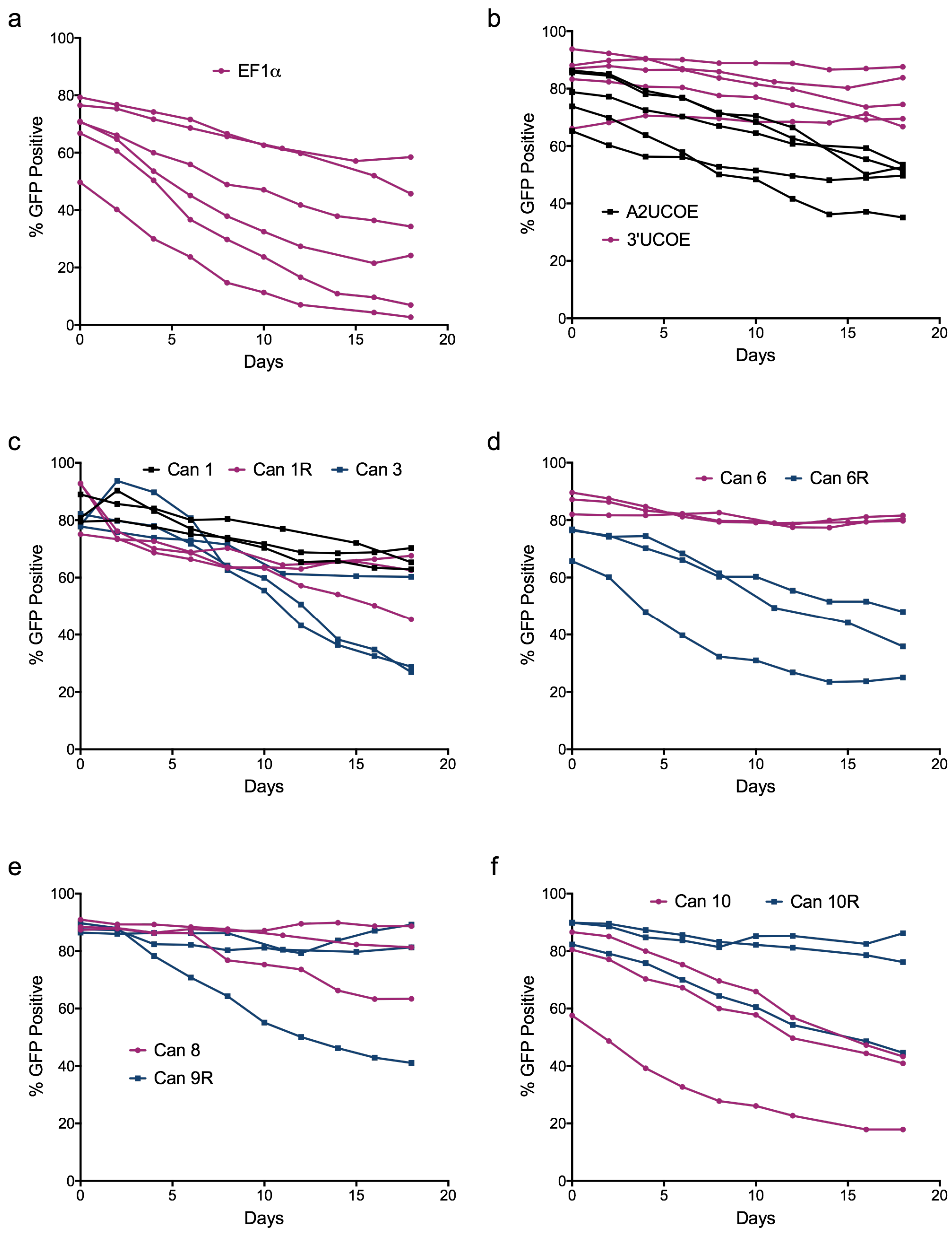

**Supplementary Figure 2. Percent GFP positive cells in the population as a function of time for UCOE candidates and controls in the P19 stable transfection screen.** After puromycin removal at day 0, cells are passaged and assayed for % GFP+ cells every 2-3 days until day 18 for the following conditions: (a) EF1a negative control, (b) A2UCOE and 3'UCOE, (c) Candidates 1, 1R, and 3, (d) Candidates 6 and 6R, (e) Candidates 8 and 9R, and (f) Candidates 10 and 10R. All replicates are shown in the same color. Data in Figure 2b is the difference between the final and initial timepoints.

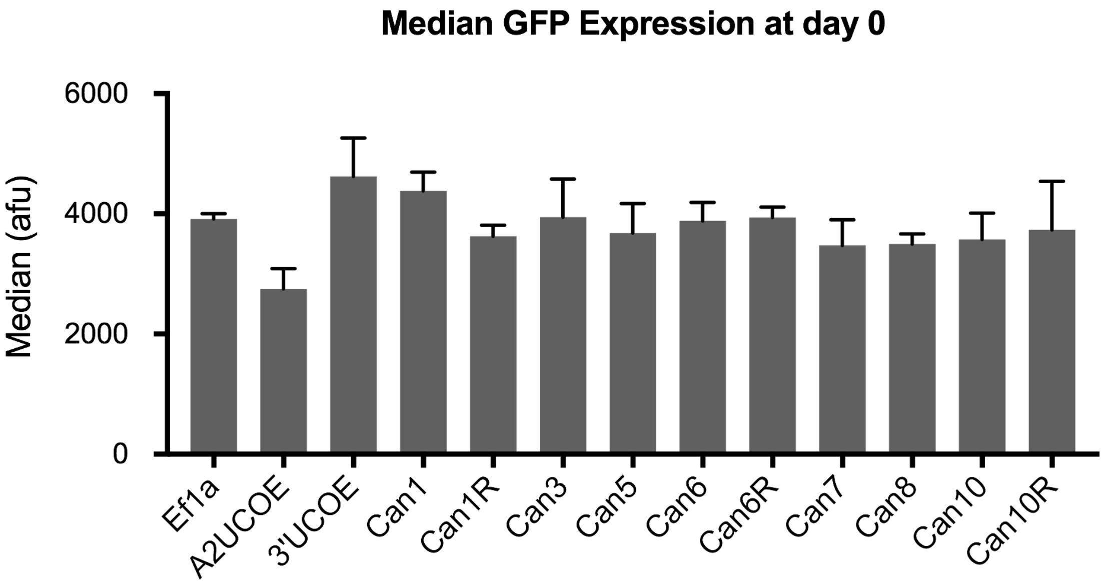

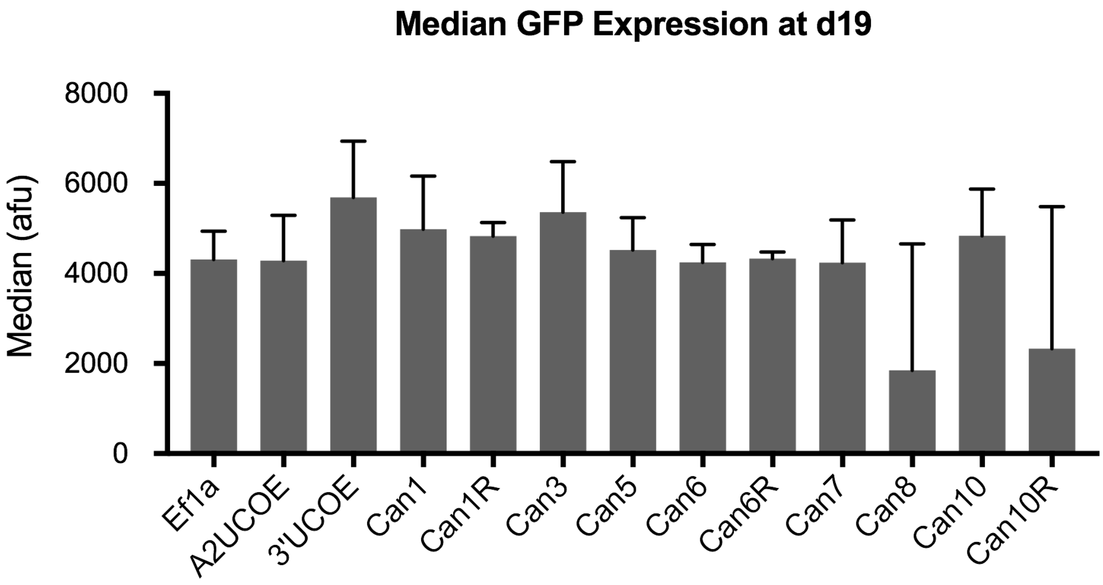

**Supplementary Figure 3. Median GFP values (in arbitrary fluorescence units) for candidate UCOEs and controls in the P19 stable transfection screen.** The median fluorescence intensity is shown for cells that are in the GFP+ gate. The data shows that median GFP values for positive controls and tested candidates are comparable to the negative (promoter-only) control at day 0 and at day 19. The unsubstantial differences in median expression demonstrate that the % GFP+ cells in the population is a more meaningful measure of silencing. Data is reported as the mean ± SD from at least three biological replicates.

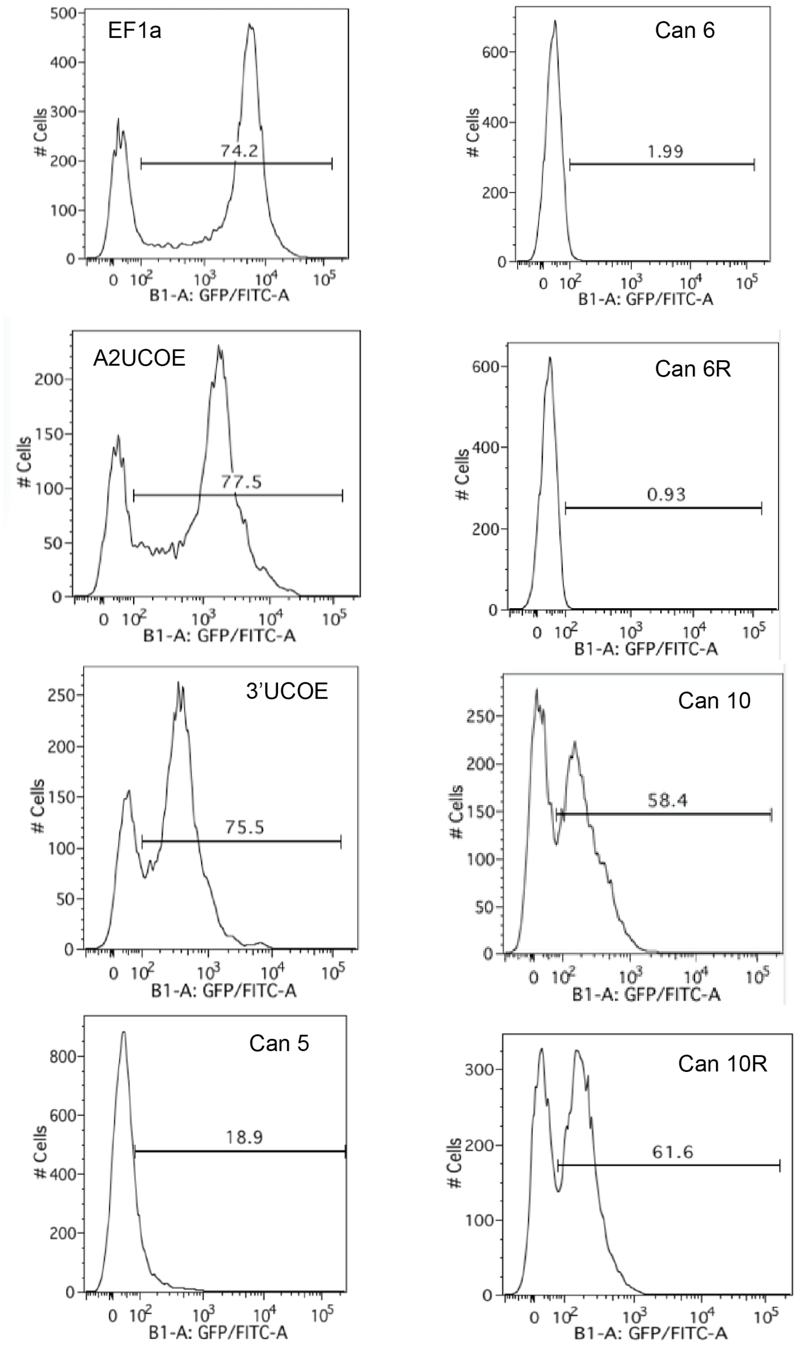

**Supplementary Figure 4. Representative histograms from flow cytometry of UCOE-GFP constructs that were stably transfected in P19s and tested for promoter activity.** Data shown here (and subsequently summarized in Figure 2c) is after 2 weeks of puromycin selection. GFP+ gate was drawn to encompass 1% of untransfected cells and then applied to all samples.

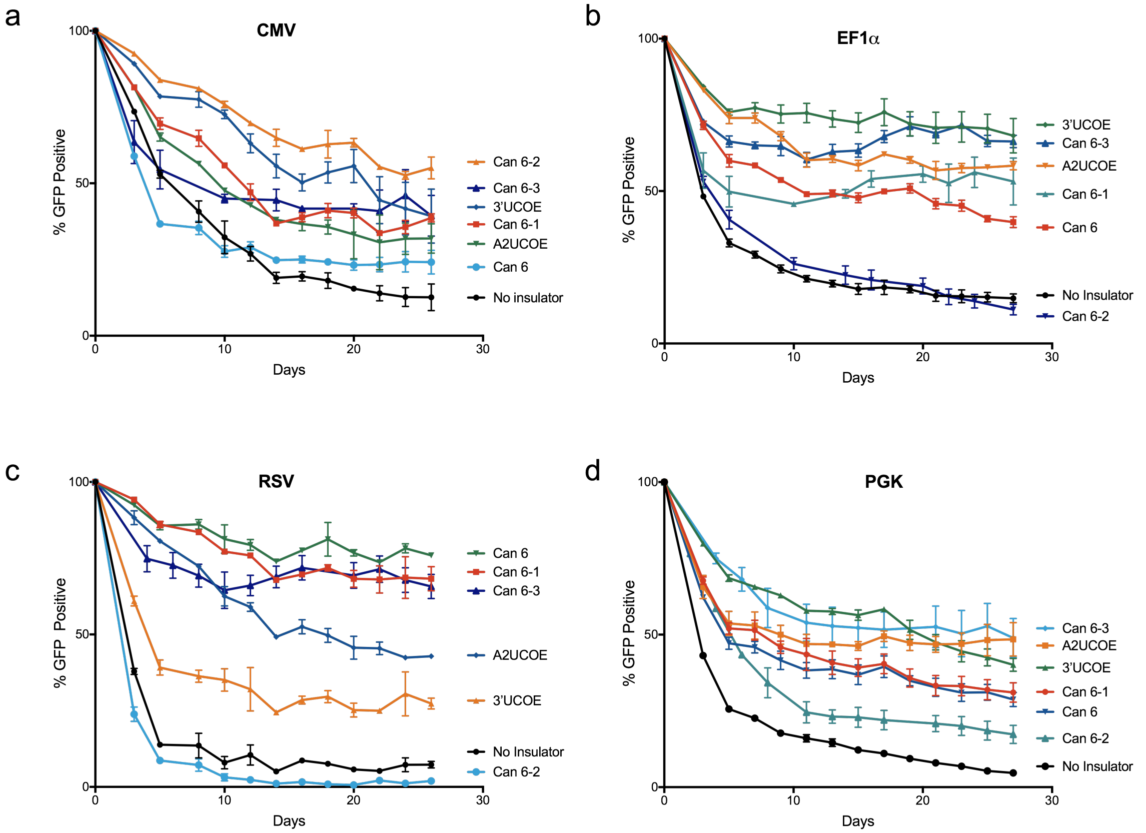

**Supplementary Figure 5. Percent GFP positive cells as a function of time for UCOE candidates and controls in the P19 lentivirus transduction assay when linked to the (a) CMV, (b) EF1a, (c) RSV, and (d) hPGK promoters.** Day 26 data is shown in Figure 4. Transduced populations were sorted into triplicate wells using FACS at day 0 (5 days after lentiviral transductions) to begin the study at 100% GFP positive cells. Expression is lost rapidly within the first 10 days, and then stabilizes around day 15. Data is reported as mean ± SD from biological triplicates.

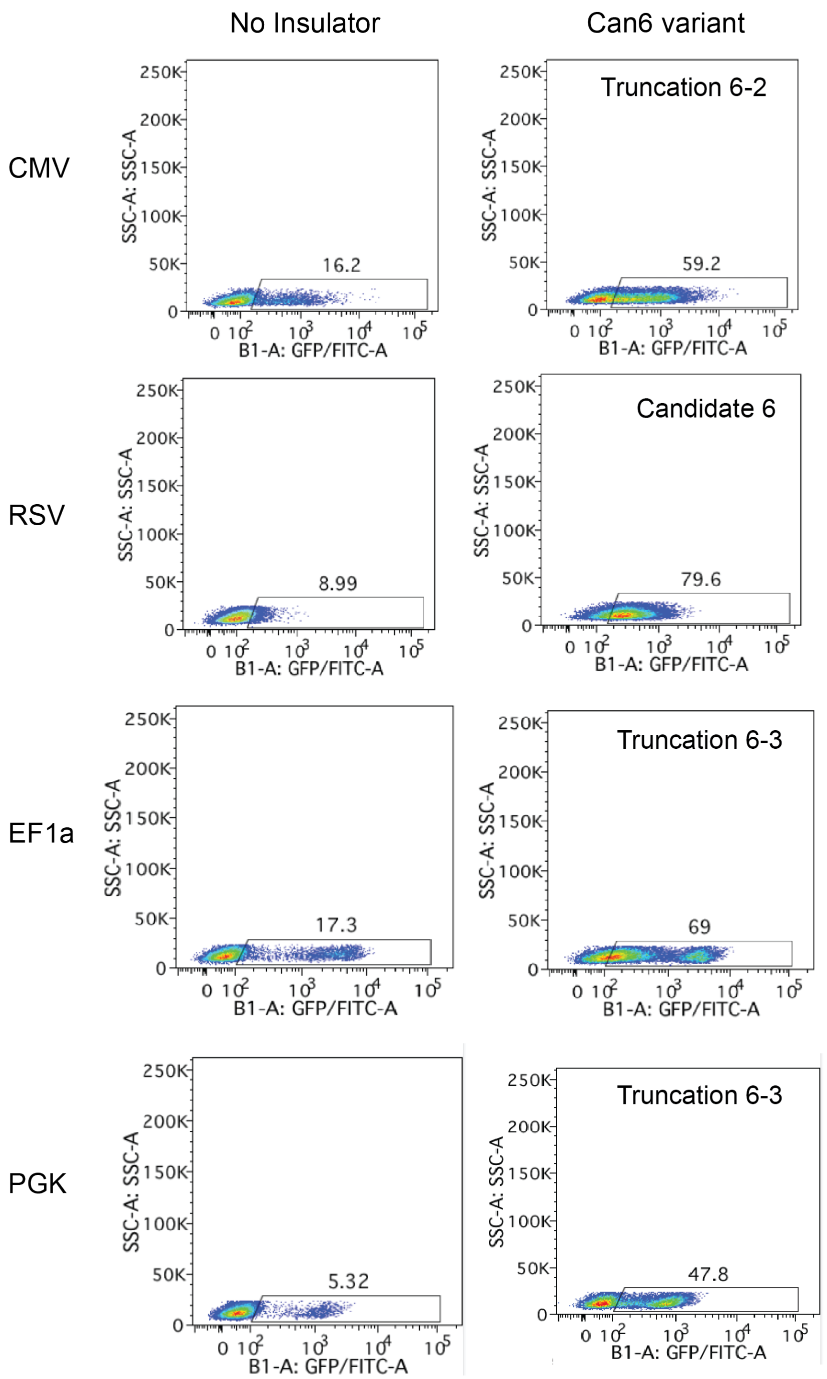

**Supplementary Figure 6. Side scatter vs. GFP for most effective Candidate 6 variants and promoter-only controls at final timepoint in the P19 lentivirus transduction assay.** Representative scatter plots are shown for each promoter-only negative control and the Candidate 6 variant (full-length or truncation) that maintained the highest %GFP+ at day 26 (CMV/RSV) or day 27 (EF1a/PGK). GFP+ gate is drawn to encompass 1% of untransduced P19 cells, and then applied to all samples. Percent GFP+ is indicated above the gate.

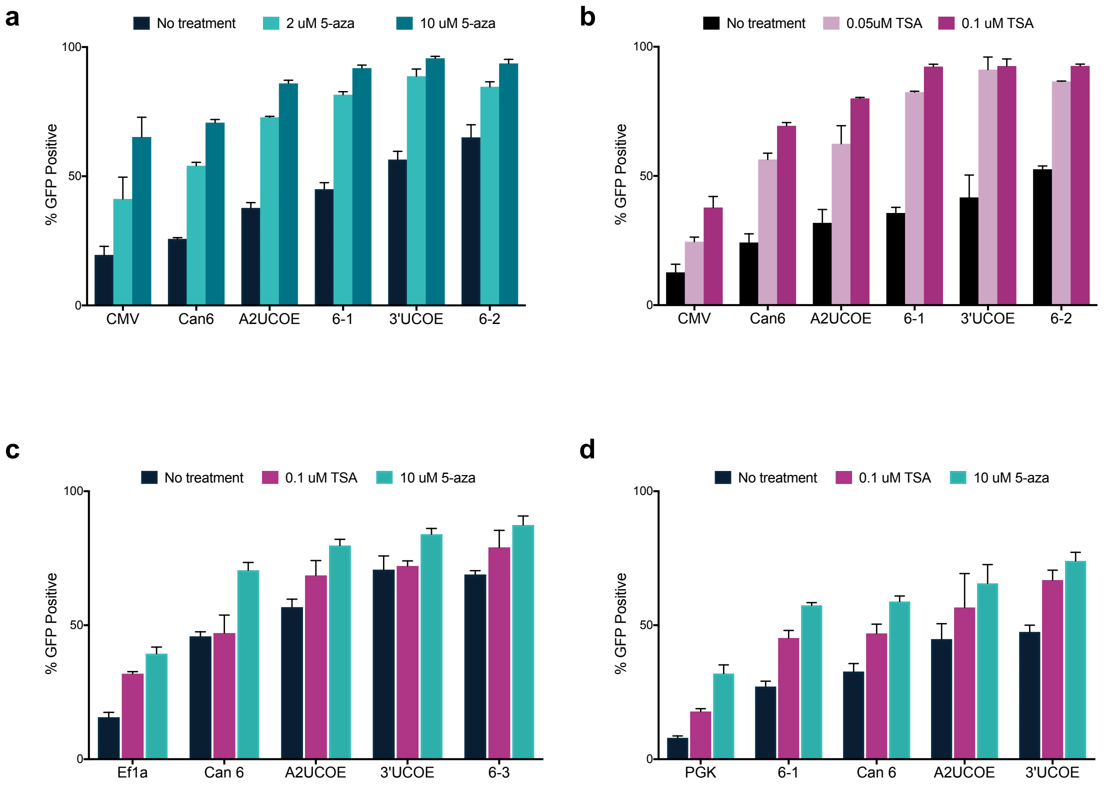

**Supplementary Figure 7. Candidate 6 resists DNA methylation and histone deacetylation.** (a) GFP expression is rescued by treatment with DNA methylation inhibitor 5-aza-cytidine (5-aza) in day 18 UCOE-CMV cells from the lentiviral silencing experiment. Cells were replica plated at day 16, specified concentrations of 5-aza were introduced 24 hours later (with exception of control), and cells were passaged and assayed via flow cytometry 24 hours after chemical introduction for % GFP+ cells. Data is reported as mean ± SD from three biological replicates. (b) GFP expression is rescued by treatment with HDAC inhibitor trichostatin A (TSA) in day 24 UCOE-CMV cells from the lentiviral silencing experiment. Cells were replica plated at day 22, specified concentrations of TSA were introduced 24 hours later (with exception of control), and cells were passaged and assayed via flow cytometry 24 hours after chemical introduction for % GFP+ cells. Data is reported as mean ± SD from three biological replicates. (c,d) UCOE candidates linked to the EF1a (c) or PGK (d) promoters demonstrate GFP expression rescue on day 21. Cells were replica plated at day 19, treated with 0.1 μM TSA or 10 μM 5-aza (with exception of control) 24 hours later, and cells were passaged and assayed via flow cytometry 24 hours after chemical introduction for % GFP+ cells. Data is reported as mean ± SD from three biological replicates.

**Supplementary Table 1. Ranked candidate UCOE regions from algorithm.** The 88 regions output by the algorithm are shown here, with chromosome position from the GRCh37/hg19 human genome assembly. Also shown is the accession number of the nearest housekeeping gene (HKG), the distance to the TSS (0 meaning the region overlaps with the TSS) and the coefficient of variation of that HKG from She et al (5). The top ranking region was the A2UCOE locus, so the next ranked candidate was labeled Candidate 1 and candidates were named sequentially.

| Candidate Rank | Chromosome | Starting Position | Ending Position | HKG Accession Number | Distance Between Candidate & HKG | CV of HKG |
|---|---|---|---|---|---|---|
| (A2UCOE) | chr7 | 26239064 | 26242980 | NM_002137 | 0 | 0.203 |
| 1 | chr6 | 160210822 | 160212252 | NM_014161 | 0 | 0.231 |
| 2 | chr1 | 245027688 | 245027958 | NM_031844 | 0 | 0.265 |
| 3 | chr6 | 160147650 | 160148658 | NM_004906 | 0 | 0.322 |
| 4 | chr16 | 25122572 | 25123049 | NM_001032391 | 0 | 0.36 |
| 5 | chr20 | 5093492 | 5094272 | NM_001009924 | 0 | 0.367 |
| 6 | chr9 | 136223261 | 136223954 | NM_017503 | 0 | 0.409 |
| 7 | chr1 | 226595430 | 226596263 | NM_001618 | 0 | 0.426 |
| 8 | chr17 | 66507666 | 66508098 | NM_212471 | 0 | 0.442 |
| 9 | chr15 | 44828479 | 44829488 | NM_003758 | 0 | 0.472 |
| 10 | chr10 | 105156061 | 105156603 | NM_032747 | 0 | 0.518 |
| 11 | chr4 | 71705461 | 71705654 | NM_002092 | 0 | 0.593 |
| 12 | chr8 | 30670056 | 30670546 | NM_001009552 | 0 | 0.634 |
| 13 | chr4 | 37891884 | 37893238 | NM_015173 | 0 | 0.773 |
| 14 | chr1 | 45476510 | 45477327 | NM_024602 | 0 | 0.845 |
| 15 | chr10 | 105156297 | 105156587 | NM_032747 | 27 | 0.518 |
| 16 | chr4 | 37892104 | 37892292 | NM_015173 | 412 | 0.773 |
| 17 | chr20 | 49548302 | 49548597 | NM_181442 | 775 | 0.317 |
| 18 | chr6 | 24720926 | 24722570 | NM_030939 | 1523 | 0.359 |
| 19 | chr13 | 76123027 | 76123946 | NM_203495 | 11019 | 0.604 |
| 20 | chr8 | 30514493 | 30515911 | NM_000637 | 19668 | 0.355 |
| 21 | chr1 | 153918934 | 153919687 | NM_020699 | 23483 | 0.315 |
| 22 | chr15 | 34659508 | 34660391 | NM_018648 | 24146 | 0.531 |
| 23 | chr4 | 2964466 | 2966496 | NM_014190 | 32664 | 0.412 |
| 24 | chr4 | 2964466 | 2966557 | NM_014190 | 32664 | 0.412 |
| 25 | chr16 | 11891244 | 11891301 | NM_015659 | 36753 | 0.414 |
| 26 | chr4 | 139936859 | 139937115 | NM_201999 | 41755 | 0.293 |

| | | | | | | |
|---|---|---|---|---|---|---|
| 27 | chr16 | 67514971 | 67515670 | NM_024519 | 47046 | 0.318 |
| 28 | chr21 | 46291815 | 46294195 | NM_006936 | 53771 | 0.266 |
| 29 | chr1 | 161008377 | 161008880 | NM_012394 | 61465 | 0.37 |
| 30 | chr11 | 65189881 | 65190103 | NM_006268 | 69430 | 0.529 |
| 31 | chr20 | 17663167 | 17663369 | NM_001011546 | 74515 | 0.731 |
| 32 | chr9 | 35814290 | 35815170 | NM_006368 | 77285 | 0.737 |
| 33 | chr17 | 37909988 | 37911027 | NM_006804 | 89534 | 0.406 |
| 34 | chr8 | 22552315 | 22553831 | NM_005775 | 119307 | 0.28 |
| 35 | chr10 | 72141721 | 72142484 | NM_021129 | 148531 | 0.432 |
| 36 | chr16 | 58035232 | 58035573 | NM_001896 | 156238 | 0.562 |
| 37 | chr1 | 23857859 | 23858006 | NM_000975 | 160262 | 0.34 |
| 38 | chr11 | 77531473 | 77532220 | NM_001293 | 182622 | 0.703 |
| 39 | chr12 | 95610899 | 95611708 | NM_018838 | 213410 | 0.489 |
| 40 | chr1 | 43389742 | 43390447 | NM_004559 | 221722 | 0.362 |
| 41 | chr10 | 72238205 | 72238357 | NM_021129 | 245015 | 0.432 |
| 42 | chr20 | 47443418 | 47445954 | NM_001037328 | 283921 | 0.29 |
| 43 | chr9 | 123639705 | 123640039 | NM_016322 | 300375 | 0.455 |
| 44 | chr22 | 19109984 | 19110404 | NM_003776 | 309631 | 0.651 |
| 45 | chr5 | 139017171 | 139017565 | NM_016480 | 311762 | 0.283 |
| 46 | chr16 | 30786630 | 30787812 | NM_024006 | 314350 | 1 |
| 47 | chr6 | 11093660 | 11094647 | NM_030969 | 336446 | 0.476 |
| 48 | chr11 | 75479737 | 75480017 | NM_001005 | 361780 | 0.422 |
| 49 | chr10 | 7452672 | 7453825 | NM_005174 | 376267 | 0.563 |
| 50 | chr11 | 18034685 | 18035793 | NM_005566 | 380142 | 0.65 |
| 51 | chr4 | 87855725 | 87856448 | NM_016245 | 401228 | 0.658 |
| 52 | chr9 | 35079517 | 35080186 | NM_007234 | 458997 | 0.456 |
| 53 | chr22 | 40742434 | 40742758 | NM_003932 | 477780 | 0.311 |
| 54 | chr6 | 52441154 | 52441900 | NM_033481 | 487895 | 0.424 |
| 55 | chr18 | 8607520 | 8610488 | NM_021074 | 492139 | 0.486 |
| 56 | chr1 | 212003870 | 212004637 | NM_018254 | 514145 | 0.445 |
| 57 | chr6 | 87861384 | 87861903 | NM_018064 | 522674 | 0.335 |
| 58 | chr2 | 153573262 | 153576806 | NM_005843 | 540756 | 0.379 |
| 59 | chr15 | 59225640 | 59226515 | NM_004492 | 703745 | 0.617 |
| 60 | chr4 | 183838726 | 183839517 | NM_152682 | 721270 | 0.528 |
| 61 | chr9 | 37034368 | 37035084 | NM_007096 | 822309 | 0.392 |
| 62 | chr1 | 221916260 | 221916494 | NM_022831 | 924860 | 0.357 |
| 63 | chr1 | 221915884 | 221916207 | NM_022831 | 925147 | 0.357 |
| 64 | chr4 | 110480710 | 110482871 | NM_000995 | 929071 | 0.419 |
| 65 | chr4 | 185570355 | 185570789 | NM_152682 | 989983 | 0.528 |
| 66 | chr14 | 91884898 | 91885777 | NM_006888 | 1010279 | 0.415 |

| | | | | | | |
|---|---|---|---|---|---|---|
| 67 | chr1 | 166808479 | 166808707 | NM_019026 | 1070320 | 0.357 |
| 68 | chr5 | 61601761 | 61601911 | NM_174889 | 1152897 | 0.362 |
| 69 | chr4 | 185747333 | 185748054 | NM_152682 | 1166961 | 0.528 |
| 70 | chr1 | 9242232 | 9242761 | NM_007262 | 1196890 | 0.363 |
| 71 | chr3 | 15468940 | 15469076 | NM_014463 | 1229071 | 0.402 |
| 72 | chr9 | 37465631 | 37467078 | NM_007096 | 1253572 | 0.392 |
| 73 | chr14 | 64107855 | 64108978 | NM_197957 | 1363840 | 0.598 |
| 74 | chr4 | 26321046 | 26324520 | NM_003102 | 1518579 | 0.436 |
| 75 | chr4 | 26321187 | 26321594 | NM_003102 | 1518720 | 0.436 |
| 76 | chr8 | 28351396 | 28352491 | NM_016127 | 1568036 | 0.397 |
| 77 | chr4 | 153700782 | 153701149 | NM_001006 | 1674978 | 0.386 |
| 78 | chr5 | 39074797 | 39075425 | NM_000997 | 1756004 | 0.45 |
| 79 | chr1 | 249151746 | 249153579 | NM_020394 | 1980351 | 0.344 |
| 80 | chr4 | 17811631 | 17812666 | NM_012161 | 2154596 | 0.421 |
| 81 | chr5 | 6711172 | 6713494 | NM_012073 | 3536538 | 0.541 |
| 82 | chr13 | 111567300 | 111568067 | NM_018011 | 4346786 | 0.246 |
| 83 | chr6 | 18264778 | 18265360 | NM_005493 | 4552982 | 0.573 |
| 84 | chr4 | 170191172 | 170193010 | NM_012403 | 5072309 | 0.38 |
| 85 | chr4 | 170192100 | 170192574 | NM_012403 | 5073237 | 0.38 |
| 86 | chr5 | 94890750 | 94891086 | NM_004365 | 5185147 | 0.813 |
| 87 | chr13 | 113343832 | 113344062 | NM_018011 | 6123318 | 0.246 |

**Supplementary Table 2. Position of algorithm output candidates versus experimental candidates in the GRCh37/hg19 human genome assembly.** Output regions from the computational search were visually inspected in UCSC Genome Browser and boundaries were drawn to result in elements around 1.5 kb while including as many features (full CpG islands, CTCF binding sites) as possible. Strand refers to +/- strand of the genome, as all candidates were drawn to be in the same 5'->3' direction as the gene with the nearest TSS to the candidate region. Regions between divergently transcribed genes are noted as divergent, along with name of the reverse complement.

| Can # | Chromo some | Algorithm start | Algorithm End | Candidate Start | Candidate End | Strand | Length | Divergent? |
|---|---|---|---|---|---|---|---|---|
| 1 | chr6 | 160210822 | 160212252 | 160210497 | 160211870 | + | 1374 | yes, 1R |
| 2 | chr1 | 245027688 | 245027958 | 245027171 | 245028685 | - | 1383 | no |
| 3 | chr6 | 160147650 | 160148658 | 160147398 | 160148705 | + | 1308 | no |
| 4 | chr16 | 25122572 | 25123049 | 25122235 | 25123617 | + | 1383 | no |
| 5 | chr20 | 5093492 | 5094272 | 5093242 | 5095057 | - | 1816 | no |
| 6 | chr9 | 136223261 | 136223954 | 136222946 | 136223954 | + | 1009* | yes, 6R |
| 7 | chr1 | 226595430 | 226596263 | 226594104 | 226596047 | + | 1944 | no |
| 8 | chr17 | 66507666 | 66508098 | 66507371 | 66509135 | + | 1765 | no |
| 9 | chr15 | 44828479 | 44829488 | 44828051 | 44829357 | + | 1307 | yes, 9R |
| 10 | chr10 | 105156061 | 105156603 | 105155621 | 105157125 | - | 1505 | yes, 10R |

**actual sequence amplified was 1002 bp as described in Supplementary Table 6*

**Supplementary Table 3. MOIs of lentiviral transductions before FACS sort.** Initial MOI based on transduction efficiency (as further described in Methods) for all data shown in Supplementary Figure 4 and Figure 4). Each set of candidates and controls are shown in order of efficacy in resisting silencing (highest to lowest %GFP+ at d26 or d27).

| Construct | MOI |
|---|---|
| **Ef1a** | |
| 3'UCOE | 0.25 |
| Can 6-3 | 0.19 |
| A2UCOE | 0.26 |
| Can6-1 | 0.29 |
| Can6 | 0.30 |
| Ef1a | 0.48 |
| Can 6-2 | 0.36 |
| **PGK** | |
| A2UCOE | 0.44 |
| Can 6-3 | 0.32 |
| 3'UCOE | 0.33 |
| Can 6-1 | 0.39 |
| Can 6 | 0.25 |
| Can 6-2 | 0.62 |
| PGK | 0.18 |
| **CMV** | |
| 6-2-CMV | 0.35 |
| A2-CMV | 0.28 |
| 6-1-CMV | 0.23 |
| 6-3-CMV | 0.57 |
| 3'UCOE-CMV | 0.28 |
| Can 6 | 0.03 |
| CMV | 0.22 |
| **RSV** | |
| Can6 | 0.38 |
| Can 6-1 | 0.12 |
| Can 6-3 | 0.48 |
| A2UCOE | 0.30 |
| 3'UCOE | 0.11 |
| RSV | 0.46 |
| 6-2-RSV | 0.53 |

**Supplementary Table 4. Plasmids used in this study.**

| Plasmid # | Description | Source/Parent |
|---|---|---|
| Stable Transfection (Screen) Plasmids | | |
| pCS3207 | pDonor for ROSA26 | Gersbach lab, Addgene #37200 |
| pCS4255 | Ef1a-EGFP | pCS3207 |
| pCS4256 | A2UCOE-Ef1a-EGFP | pCS4255 |
| pCS4257 | 3'UCOE-Ef1a-EGFP | pCS4255 |
| pCS4258 | Candidate1-Ef1a-EGFP | pCS4257 |
| pCS4259 | Candidate1R-Ef1a-EGFP | pCS4257 |
| pCS4260 | Can3-Ef1a-EGFP | pCS4257 |
| pCS4261 | Can6-Ef1a-EGFP | pCS4257 |
| pCS4262 | Can6(opp)-EF1a-EGFP | pCS4257 |
| pCS4263 | Can8-Ef1a-EGFP | pCS4257 |
| pCS4264 | Can9R-Ef1a-EGFP | pCS4257 |
| pCS4265 | A2UCOE-EGFP | pCS4255 |
| pCS4266 | 3'UCOE-EGFP | pCS4255 |
| pCS4267 | Can1-EGFP | pCS4266 |
| pCS4268 | Can1(opp)-EGFP | pCS4266 |
| pCS4269 | Can3-EGFP | pCS4266 |
| pCS4270 | Can6-EGFP | pCS4266 |
| pCS4271 | Can6(opp)-EGFP | pCS4266 |
| pCS4272 | Can5-Ef1a-EGFP | pCS4266 |
| pCS4273 | Can7-Ef1a-EGFP | pCS4266 |
| pCS4274 | Can10-Ef1a-EGFP | pCS4266 |
| pCS4275 | Can10R-Ef1a-EGFP | pCS4266 |
| Lenti plasmids | | |
| pCS3799 | pLenti donor | Xiang et al (6) |
| pCS3800 | pMO86 HIV-1 Gag packaging plasmid | Xiang et al (6) |
| pCS3801 | pMO87 VSV g envelope protein plasmid | Xiang et al (6) |
| pCS4276 | pKL5-with ef1a-EGFP | pCS3799 |
| pCS4277 | pKL5-with Can6-ef1a-EGFP | pCS3799 |
| pCS4278 | pKL5 with Can12R-ef1a-EGFP | pCS3799 |
| pCS4279 | pKL5 with A2UCOE-Ef1a-EGFP | pCS3799 |
| pCS4280 | pKL5 with 3'ucoe-ef1a-EGFP | pCS3799 |
| pCS4281 | Can6-1-ef1a-EGFP | pCS4278 |
| pCS4282 | Can6-2-ef1a-EGFP | pCS4278 |
| pCS4283 | 6-3-ef1a-EGFP | pCS4278 |
| pCS4284 | pKL5 with CMV-EGFP | pCS3799 |
| pCS4285 | Can6-1-CMV-EGFP | pCS4284 |

| | | |
|---|---|---|
| pCS4286 | Can6-2-CMV-EGFP | pCS4284 |
| pCS4287 | A2UCOE-CMV-EGFP | pCS4284 |
| pCS4288 | 3'UCOE-CMV-EGFP | pCS4284 |
| pCS4289 | Can6-CMV-EGFP | pCS4284 |
| pCS4290 | Can 6-3-CMV | pCS4284 |
| pCS4291 | pKL5 with PGK-EGFP | pCS3799 |
| pCS4292 | Can6-1-PGK-EGFP | pCS4291 |
| pCS4293 | Can6-2-PGK-EGFP | pCS4291 |
| pCS4294 | A2UCOE-PGK-EGFP | pCS4291 |
| pCS4295 | 3'UCOE-PGK-EGFP | pCS4291 |
| pCS4296 | Can6-PGK-EGFP | pCS4291 |
| pCS4297 | Can 6-3-PGK | pCS4292 |
| pCS4298 | RSV-EGFP | pCS4291 |
| pCS4299 | Can6-1-RSV-EGFP | pCS4298 |
| pCS4300 | 3'UCOE-RSV-EGFP | pCS4295 |
| pCS4301 | Can6-RSV-EGFP | pCS4296 |
| pCS4302 | A2UCOE-RSV-EGFP | pCS4294 |
| pCS4303 | Can6-2-RSV-EGFP | pCS4298 |
| pCS4304 | Can 6-3-RSV-EGFP | pCS4303 |

**Supplementary Table 5. Primers used in this study.**

| | |
|---|---|
| Can1_fwd_Sal1 | actaagaGTCGACCCATCTTGACGGCAGCGATA |
| Can1_rev_Nhe1 | tagttctGCTAGCCGCTGAGACGATCTCGGAAA |
| Can1R_fwd_Sal1 | actaagaGTCGACCGCTGAGACGATCTCGGAAA |
| Can1R_rev_Nhe1 | tagttctGCTAGCCCATCTTGACGGCAGCGATA |
| Can2_fwd_Sal1 | actaagaGTCGACTCACCCTCACGGTTAGCTACT |
| Can2_rev_Nhe1 | tagttctGCTAGCCAACGTACAACGCAGCACTC |
| Can3_fwd_Sal1 | actaagaGTCGACCAGACCGATCTGATTCACTGG |
| Can3_rev_Nhe1 | tagttctGCTAGCCGGTCGCATAGGCCGAG |
| Can4_fwd_Sal1 | actaagaGTCGACACTTTTCCACACACTACTTCCCTC |
| Can4_rev_Nhe1 | tagttctGCTAGCTCTGTCTTTCCAGCAGCGTT |
| Can6_fwd_Sal1 | actaagaGTCGACGCACACGACCACAATTCCAC |
| Can6_rev_Nhe1 | tagttctGCTAGCGACCACCTACGGGTTCTTGG |
| Can6R_fwd_Sal1 | actaagaGTCGACGACCACCTACGGGTTCTTGG |
| Can6R_rev_Nhe1 | tagttctGCTAGCGCACACGACCACAATTCCAC |
| Can8_fwd_Sal1 | actaagaGTCGACAAGCACACGGCCCTAGAAAT |
| Can8_rev_Nhe1 | tagttctGCTAGCTGGAGAGGAAAACTACCGGC |
| Can9_fwd_Sal1 | actaagaGTCGACGTCCTGCCCACGTATCTACC |
| Can9_rev_Nhe1 | tagttctGCTAGCCTAGCGAGGAGTTAGCACGG |
| Can9R_fwd_Sal1 | actaagaGTCGACCTAGCGAGGAGTTAGCACGG |
| Can9R_rev_Nhe1 | tagttctGCTAGCGTCCTGCCCACGTATCTACC |
| Can5_fwd_Sal1 | actaagaGTCGACCAAGTTCACTGTGTGCTGTGTATT |
| Can5_rev_Nhe1 | tagttctGCTAGCGTCTTCGTTGCCAACAGGCT |
| Can7_fwd_Sal1 | actaagaGTCGAC GAGGGGTTGGGGGTAAAATTAGT |
| Can7_rev_Nhe1 | tagttctGCTAGC AGGTTCCTTAGTGGGCAACA |
| Can10_fwd_Sal1 | actaagaGTCGAC AGCAGGGAAAGCGAGAGAAC |
| Can10_rev_Nhe1 | tagttctGCTAGC AAAGGCCTTCCCACTGATCG |
| Can10R_fwd_Sal1 | actaagaGTCGAC AAAGGCCTTCCCACTGATCG |
| Can10R_rev_Nhe1 | tagttctGCTAGC AGCAGGGAAAGCGAGAGAAC |
| Can6_213F_fwd (for 6-1) | actaagaGTCGAC TTCAAAGTGCAGGGCAGACA |
| Can6_585F_fwd (for 6-2) | actaagaGTCGAC TTCTGCGAGCGGCTTCC |
| Can6_874R_rev (for 6-3) | tagttctGCTAGC TTCCCTCTCCTCCCCTGATC |

**Supplementary Table 6. SRF-UCOE and promoter sequences.**

| | |
|---|---|
| **Candidate 6 (SRF-UCOE) – 1002 bp** | GCACACGACCACAATTCCACTGAAAGCATTTTAATACGG<br>AACTTGTCACTCCCAGGGAGCCTCCGCTCAGCCGGCAGTT<br>GGTTCATTTCAATCCCCACGACAACCCTTCAAAGTGCAGG<br>GCAGACAGCAGGTGGCTCTGCCCAGGCGCCTGGATCACA<br>GCCCGGCCTGCAGCCCTCACCTGGGCGCGGGGAGACCCT<br>GAGGACGCTCCTCCAGGCGGCGCTGGCCGGGGCCTGCGG<br>ACACGGACGGGCGGGCTGAGCTCCGGGACCCCTCCCCGC<br>GCCCCGCACCCCGCACCCCGCACCCCGCACCCCGCACCC<br>GGCGCTCACCCGTCCCAGCCCCGCCGCCCGCAGCCCCAG<br>CTGCAACGCAGCCACCGCCGCCATCGCACCCGGCCCCGC<br>GGGCGCTTCCGGGACGCAGGAGGCATCTGCATCCGGGGC<br>GCCGCTGAGTCCCGCCCAGAGCCCCGCCCCCGGCTCCAG<br>GTTCTGCGAGCGGCTTCCGCCGGGCTGCTCCGCGGGCGCG<br>TCGGCCATGAGCGAGTTGCCGGGCGACGTGCGGGCGTTT<br>CTGCGGGAGCACCCGAGCCTGCGGCTCCAGACGGACGCC<br>CGCAAGGTTCGCAGCGCGGGAGGGGAACGGAGTGGCGG<br>AGAAGGGCGCAGTTGGGATGAGGGGCTGAGGGGAGGGC<br>AGGGGAGAGGAGAGGGCAGGGGAGAGGGGAGAGGGGA<br>GAGCAGGAGAGAGGGGAAGGCAGGGGAGAGGGCGCGGC<br>GGGATCAGGGGAGGAGAGGGAAGGGGGCGCGGCAGGAG<br>GGGGCACCAGGGAGCGGAGCCCTGGCCCTCCTGACGTCC<br>TGCCCGCCCACGCGTCCGCAGGTGAGGTGCATCCTGACA<br>GGTCACGAGCTGCCCTGCCGCCTGCCGGAGCTCCAGGTCT<br>ACACCCGCGGCAAAAAGTACCAGCGGCTGGTCCGCGCCT<br>CCCCGGCCTTCGACTATGCAGAGTTCGAGCCGCACATCGT<br>GCCCAGCACCAAGAACCCGTAGGTGGTC |
| **Truncation 6-1 (896bp)** | TTCAAAGTGCAGGGCAGACAGCAGGTGGCTCTGCCCAGG<br>CGCCTGGATCACAGCCCGGCCTGCAGCCCTCACCTGGGC<br>GCGGGGAGACCCTGAGGACGCTCCTCCAGGCGGCGCTGG<br>CCGGGGCCTGCGGACACGGACGGGCGGGCTGAGCTCCGG<br>GACCCCTCCCCGCGCCCCGCACCCCGCACCCCGCACCCCG<br>CACCCCGCACCCGGCGCTCACCCGTCCCAGCCCCGCCGCC<br>CGCAGCCCCAGCTGCAACGCAGCCACCGCCGCCATCGCA<br>CCCGGCCCCGCGGGCGCTTCCGGGACGCAGGAGGCATCT<br>GCATCCGGGGCGCCGCTGAGTCCCGCCCAGAGCCCCGCC<br>CCCGGCTCCAGGTTCTGCGAGCGGCTTCCGCCGGGCTGCT<br>CCGCGGGCGCGTCGGCCATGAGCGAGTTGCCGGGCGACG<br>TGCGGGCGTTTCTGCGGGAGCACCCGAGCCTGCGGCTCC<br>AGACGGACGCCCGCAAGGTTCGCAGCGCGGGAGGGGAA<br>CGGAGTGGCGGAGAAGGGCGCAGTTGGGATGAGGGGCT<br>GAGGGGAGGGCAGGGGAGAGGAGAGGGCAGGGGAGAG<br>GGGAGAGGGGAGAGCAGGAGAGAGGGGAAGGCAGGGG<br>AGAGGGCGCGGCGGGATCAGGGGAGGAGAGGGAAGGGG<br>GCGCGGCAGGAGGGGGCACCAGGGAGCGGAGCCCTGGC<br>CCTCCTGACGTCCTGCCCGCCCACGCGTCCGCAGGTGAGG<br>TGCATCCTGACAGGTCACGAGCTGCCCTGCCGCCTGCCGG<br>AGCTCCAGGTCTACACCCGCGGCAAAAAGTACCAGCGGC<br>TGGTCCGCGCCTCCCCGGCCTTCGACTATGCAGAGTTCGA |

| | |
|---|---|
| | GCCGCACATCGTGCCCAGCACCAAGAACCCGTAGGTGGT C |
| **Truncation 6-2 (524bp)** | AGCGGCTTCCGCCGGGCTGCTCCGCGGGCGCGTCGGCCA TGAGCGAGTTGCCGGGCGACGTGCGGGCGTTTCTGCGGG AGCACCCGAGCCTGCGGCTCCAGACGGACGCCCGCAAGG TTCGCAGCGCGGGAGGGGAACGGAGTGGCGGAGAAGGG CGCAGTTGGGATGAGGGGCTGAGGGGAGGGCAGGGGAG AGGAGAGGGCAGGGGAGAGGGGAGAGGGGAGAGCAGG AGAGAGGGGAAGGCAGGGGAGAGGGCGCGGCGGGATCA GGGGAGGAGAGGGAAGGGGGCGCGGCAGGAGGGGGCAC CAGGGAGCGGAGCCCTGGCCCTCCTGACGTCCTGCCCGC CCACGCGTCCGCAGGTGAGGTGCATCCTGACAGGTCACG AGCTGCCCTGCCGCCTGCCGGAGCTCCAGGTCTACACCCG CGGCAAAAAGTACCAGCGGCTGGTCCGCGCCTCCCCGGC CTTCGACTATGCAGAGTTCGAGCCGCACATCGTGCCCAGC ACCAAGAACCCGTAGGTGGTC |
| **Truncation 6-3 (761bp)** | GCACACGACCACAATTCCACTGAAAGCATTTTAATACGG AACTTGTCACTCCCAGGGAGCCTCCGCTCAGCCGGCAGTT GGTTCATTTCAATCCCCACGACAACCCTTCAAAGTGCAGG GCAGACAGCAGGTGGCTCTGCCCAGGCGCCTGGATCACA GCCCGGCCTGCAGCCCTCACCTGGGCGCGGGGAGACCCT GAGGACGCTCCTCCAGGCGGCGCTGGCCGGGGCCTGCGG ACACGGACGGGCGGGCTGAGCTCCGGGACCCCTCCCCGC GCCCCGCACCCCGCACCCCGCACCCCGCACCCCGCACCC GGCGCTCACCCGTCCCAGCCCCGCCGCCCGCAGCCCCAG CTGCAACGCAGCCACCGCCGCCATCGCACCCGGCCCCGC GGGCGCTTCCGGGACGCAGGAGGCATCTGCATCCGGGGC GCCGCTGAGTCCCGCCCAGAGCCCCGCCCCCGGCTCCAG GTTCTGCGAGCGGCTTCCGCCGGGCTGCTCCGCGGGCGCG TCGGCCATGAGCGAGTTGCCGGGCGACGTGCGGGCGTTT CTGCGGGAGCACCCGAGCCTGCGGCTCCAGACGGACGCC CGCAAGGTTCGCAGCGCGGGAGGGGAACGGAGTGGCGG AGAAGGGCGCAGTTGGGATGAGGGGCTGAGGGGAGGGC AGGGGAGAGGAGAGGGCAGGGGAGAGGGGAGAGGGGA GAGCAGGAGAGAGGGGAAGGCAGGGGAGAGGGCGCGGC GGGATCAGGGGAGGAGAGGGAA |
| **Promoter- RSV** | AATGTAGTCTTATGCAATACTCTTGTAGTCTTGCAACATG GTAACGATGAGTTAGCAACATGCCTTACAAGGAGAGAAA AAGCACCGTGCATGCCGATTGGTGGAAGTAAGGTGGTAC GATCGTGCCTTATTAGGAAGGCAACAGACGGGTCTGACA TGGATTGGACGAACCACTGAATTCCGCATTGCAGAGATA TTGTATTTAAGTGCCTAGCTCGATACAATAAACGCCATTT GACCATTCACCACATTGGTGTGCACC |
| **Promoter- CMV** | GTTGACATTGATTATTGACTAGTTATTAATAGTAATCAAT TACGGGGTCATTAGTTCATAGCCCATATATGGAGTTCCGC GTTACATAACTTACGGTAAATGGCCCGCCTGGCTGACCGC CCAACGACCCCCGCCCATTGACGTCAATAATGACGTATGT TCCCATAGTAACGCCAATAGGGACTTTCCATTGACGTCAA TGGGTGGAGTATTTACGGTAAACTGCCCACTTGGCAGTAC |

| | |
|---|---|
| | ATCAAGTGTATCATATGCCAAGTACGCCCCCTATTGACGT CAATGACGGTAAATGGCCCGCCTGGCATTATGCCCAGTA CATGACCTTATGGGACTTTCCTACTTGGCAGTACATCTAC GTATTAGTCATCGCTATTACCATGGTGATGCGGTTTTGGC AGTACATCAATGGGCGTGGATAGCGGTTTGACTCACGGG GATTTCCAAGTCTCCACCCCATTGACGTCAATGGGAGTTT GTTTTGGCACCAAAATCAACGGGACTTTCCAAAATGTCGT AACAACTCCGCCCCATTGACGCAAATGGGCGGTAGGCGT GTACGGTGGGAGGTCTATATAAGCAGAGCTCT |
| **Promoter- Ef1a** | GATCTGCGATCGCTCCGGTGCCCGTCAGTGGGCAGAGCG CACATCGCCCACAGTCCCCGAGAAGTTGGGGGGAGGGGT CGGCAATTGAACCGGTGCCTAGAGAAGGTGGCGCGGGGT AAACTGGGAAAGTGATGTCGTGTACTGGCTCCGCCTTTTT CCCGAGGGTGGGGGAGAACCGTATATAAGTGCAGTAGTC GCCGTGAACGTTCTTTTTCGCAACGGGTTTGCCGCCAGAA CACAGCTGAAGCTTCGAGGGGCTCGCATCTCTCCTTCACG CGCCCGCCGCCCTACCTGAGGCCGCCATCCACGCCGGTTG AGTCGCGTTCTGCCGCCTCCCGCCTGTGGTGCCTCCTGAA CTGCGTCCGCCGTCTAGGTAAGTTTAAAGCTCAGGTCGAG ACCGGGCCTTTGTCCGGCGCTCCCTTGGAGCCTACCTAGA CTCAGCCGGCTCTCCACGCTTTGCCTGACCCTGCTTGCTC ACCTCTACGTCTTTGTTTCGTTTTCTGTTCTGCGCCGTTAC AGATCCAAGCTGTGACCGGCGCCTAC |
| **Promoter- PGK** | CCACGGGGTTGGGGTTGCGCCTTTTCCAAGGCAGCCCTGG GTTTGCGCAGGGACGCGGCTGCTCTGGGCGTGGTTCCGG GAAACGCAGCGGCGCCGACCCTGGGTCTCGCACATTCTTC ACGTCCGTTCGCAGCGTCACCCGGATCTTCGCCGCTACCC TTGTGGGCCCCCCGGCGACGCTTCCTGCTCCGCCCCTAAG TCGGGAAGGTTCCTTGCGGTTCGCGGCGTGCCGGACGTG ACAAACGGAAGCCGCACGTCTCACTAGTACCCTCGCAGA CGGACAGCGCCAGGGAGCAATGGCAGCGCGCCGACCGCG ATGGGCTGTGGCCAATAGCGGCTGCTCAGCGGGCGCGCC GAGAGCAGCGGCCGGGAAGGGGCGGTGCGGGAGGCGGG GTGTGGGGCGGTAGTGTGGGCCCTGTTCCTGCCCGCGCGG TGTTCCGCATTCTGCAAGCCTCCGGAGCGCACGTCGGCAG TCGGCTCCCTCGTTGACCGAATCACCGACCTCTCTCCCCA GG |

**Supplementary Material References**

1. Guttman, M., Garber, M., Levin, J.Z., Donaghey, J., Robinson, J., Adiconis, X., Fan, L., Koziol, M.J., Gnirke, A., Nusbaum, C. *et al.* (2010) Ab initio reconstruction of cell type-specific transcriptomes in mouse reveals the conserved multi-exonic structure of lincRNAs. *Nat Biotechnol*, **28**, 503-510.
2. Hsu, F., Kent, W.J., Clawson, H., Kuhn, R.M., Diekhans, M. and Haussler, D. (2006) The UCSC Known Genes. *Bioinformatics*, **22**, 1036-1046.
3. Consortium, E.P. (2012) An integrated encyclopedia of DNA elements in the human genome. *Nature*, **489**, 57-74.
4. Wang, J., Zhuang, J., Iyer, S., Lin, X.Y., Greven, M.C., Kim, B.H., Moore, J., Pierce, B.G., Dong, X., Virgil, D. *et al.* (2013) Factorbook.org: a Wiki-based database for transcription factor-binding data generated by the ENCODE consortium. *Nucleic Acids Res*, **41**, D171-176.
5. She, X., Rohl, C.A., Castle, J.C., Kulkarni, A.V., Johnson, J.M. and Chen, R. (2009) Definition, conservation and epigenetics of housekeeping and tissue-enriched genes. *BMC Genomics*, **10**, 269.
6. Xiang, J.S.K., M.; Dykstra, P.; Hinks, M.; McKeague, M.; Smolke, C.D. (2019) Massively Parallel RNA Device Engineering in Mammalian Cells with RNA-Seq and FACS-Seq. *(under review)*.
